# One night of sleep deprivation reduces spatially patterned human brain clearance: implications for simulated tau propagation

**DOI:** 10.1101/2025.06.03.657596

**Authors:** Georgia S. Brennan, Travis B. Thompson, Hadrien Oliveri, Vegard Vinje, Geir Ringstad, Per Kristian Eide, Alain Goriely, Marie E. Rognes

## Abstract

Neurotoxic protein fragments such as amyloid-beta and tau accumulate in characteristic staging patterns in Alzheimer’s disease. The brain clears such substances via multiple systems, including the extracellular–extravascular route known as glymphatic clearance, but how such clearance is distributed across the brain, how this distribution is affected by sleep, and its implications for neurodegenerative disease progression remain unclear. Using Bayesian inference to fit a kinetic model of enhancement and clearance to longitudinal CSF tracer-enhanced MRI, we estimate CSF tracer clearance rates for every connectome region in 21 subjects. We find that clearance is spatially patterned across the living human brain and consistent between subjects. Clearance rates are amongst the highest in the entorhinal cortex and decrease across regions in Braak stage order. One night of sleep deprivation reduces mean clearance levels while leaving the spatial pattern largely unchanged. To study how mean clearance level and spatial pattern affect tau propagation in silico, we use the CSF tracer clearance rates as a proxy for regional tau clearance in a computational model of protein accumulation, spread, and regional damage. Across subject-specific simulations, clearance level and spatial pattern have distinct effects on tau propagation. The clearance level determines the timescale of simulated pathology, while the spatial pattern shapes its accumulation and onset. These results identify CSF tracer clearance as a spatially organised property of the human brain that is reduced by sleep deprivation, and suggest that its overall level and regional organisation may independently shape the timing and spatial progression of neurodegenerative proteinopathy.

**SIGNIFICANCE STATEMENT:** Regional brain clearance in the living human brain is strongly organised and broadly consistent between subjects: it is among the fastest in the entorhinal cortex, where tau pathology first appears, and declines across the Braak sequence. One night of sleep deprivation lowered the overall clearance rate by approximately one third while largely preserving which regions cleared fastest. Applying these regional clearance rates as initial conditions in a connectome model of simulated protein propagation, the overall rate determined when simulated tau accumulation occurred, whereas the regional organisation delayed its onset and influenced its anatomical distribution. These modelled effects describe mechanistic implications of the measured clearance maps and are not observations of clinical disease progression.

## Introduction

The brain represents just 2% of the average human’s body weight, but consumes approximately 20% of the body’s energy^1^. The by-products of its metabolism include protein fragments such as misfolded amyloid-β (Aβ), tau proteins (τ) and α-synuclein, which are implicated in the onset and progression of neurodegenerative diseases^2^. Lacking classical lymphatic vessels^3,4^, the brain removes such metabolic waste and other solutes through several other clearance pathways^5^. These include transport across the blood-brain barrier, intracellular degradation and autophagy, and glymphatic – extracellular and extravascular – clearance mediated by interstitial fluid (ISF) and cerebrospinal fluid (CSF) flow^6,7^. A growing body of evidence suggests that the glymphatic pathway plays an important role in maintaining healthy brain clearance^8^. Impaired glymphatic function reduces Aβ^6,9^ and τ^10,11^ clearance in mice, with early results indicating that the same may hold also for α-synuclein^12,13^.

In humans, intrathecal contrast-enhanced glymphatic magnetic resonance imaging (gMRI), a longitudinal neuroimaging protocol using an MRI contrast agent as a CSF tracer, is now established as a technique to study brain solute transport^7,14–17^. These studies have revealed brain-wide tracer transport with marked regional differences in both enrichment and clearance. Tracer distribution is also sensitive to physiological state, as vascular and intracranial pulsatility are closely coupled to CSF flow, ISF–CSF exchange, and flow-enhanced transport^18–20^. Moreover, sleep enhances solute movement and clearance^21,22^, in part through slow vasomotion^20,23^, while sleep deprivation delays brain clearance^24^. Yet, the downstream implications of how glymphatic clearance and its physiological variations may modulate neurodegenerative disease remain hard to quantify.

Mathematical modelling provides a complementary route to study the influence of brain solute clearance on neurodegenerative disease development over longer timescales. Models of Aβ and τ aggregation, propagation, and clearance dynamics across the human brain connectome^25–28^ demonstrate that enhanced clearance can delay features of Alzheimer’s disease (AD) onset and progression. In particular, spatially-varying clearance rates can alter the simulated progression of protein pathology and contribute to the manifestation of distinct AD subtypes^26^.

In this study, we address three key open questions. First, what is the spatial organisation of CSF tracer clearance in the living human brain? Second, does one night of sleep deprivation alter its overall level, its regional pattern, or both? Third, how do such gMRI-informed spatial clearance maps, their overall level and regional patterns, influence simulated τ propagation across the brain connectome? Specifically, using Bayesian techniques, we estimate and analyse regional clearance rates from previously collected human gMRI data^24^, including both reference and sleep-deprived subjects, to generate subject-specific clearance maps across the connectome. We then use these clearance maps to initialise a connectome-based model of protein accumulation, spread, and regional damage, to simulate AD progression in each subject. We emphasise that these imaging-based clearance maps represent non-AD reference subjects, and thus we consider how physiological clearance characteristics may protect against τ propagation rather than aiming to replicate AD trajectories.

We find that human CSF tracer clearance displays spatial patterns that are consistent across subjects, with clearance rates among the highest in the entorhinal cortex, where τ pathology first appears, and decreasing across the Braak hierarchy. One night of sleep deprivation lowers clearance widely across the brain, at every Braak stage and in every cortical lobe, while largely preserving this pattern. Clearance level and spatial pattern have distinct effects on simulated tauopathy. The average clearance rate sets the timescale of progression, with a twofold range between subjects corresponding to a 17.5-year spread in time to widespread pathology. The spatial pattern delays the first appearance of simulated pathology by about three years relative to spatially uniform clearance at the same mean rate, and contributes to which regions are affected first. Sleep deprivation shifts simulated progression earlier by lowering the clearance level, and thus reducing the protection afforded by high clearance in the entorhinal cortex. Together, these findings identify the spatial structure and overall level of human brain clearance as potential modulators of neurodegenerative disease progression and quantify how sleep deprivation perturbs this system.

## Results

In a previously reported study^24^, MRI in two patient cohorts followed an intrathecally injected CSF tracer as it spread through the brain and was cleared over the following days. From those images, we estimate, for each subject and for each region of the connectome, the rate at which tracer left the tissue. We refer to each subject’s set of rates as their *clearance map*. This map has two key properties: its *level*, defined as the mean clearance rate, and its spatial *pattern*, defined by how clearance rates are distributed across regions. Next, we impose these maps on a computational model of τ propagation across the connectome, and ask how level and pattern each affect the simulated disease progression. Both the clearance maps and the simulations are compared between the reference and sleep-deprived cohorts.

### Regional clearance varies over the connectome with consistent spatial patterns across subjects

To map from volumetric gMRI data describing contrast-induced T1 signal intensity at multiple time points to a *clearance map* across the connectome, we fit a two-compartment model to the full time course for each region of interest (ROI) in *n* = 21 subjects. The subjects comprise a reference cohort (*n* = 14), who slept normally on the night after injection, and a sleep-deprived cohort (*n* = 7), who were kept awake through that night^24^. The two-compartment model jointly estimates tracer enhancement rate *k*_in,*i*_, the clearance rate *k*_out,*i*_, and the peak amplitude *A*_*i*_ in each ROI *i*, using hierarchical Bayesian inference with cohort-specific means and partial pooling across all subjects and reporting on posterior medians (Figure 1, Methods). The clearance rates *{k*_out,*i*_*}* _*i*_ define a subject-specific map of clearance, in units of per hour, across 42 regions: the 34 cortical regions of the Desikan–Killiany parcellation, 7 subcortical structures from the FreeSurfer subcortical segmentation, and the brain stem^29,30^.

**Figure 1.**
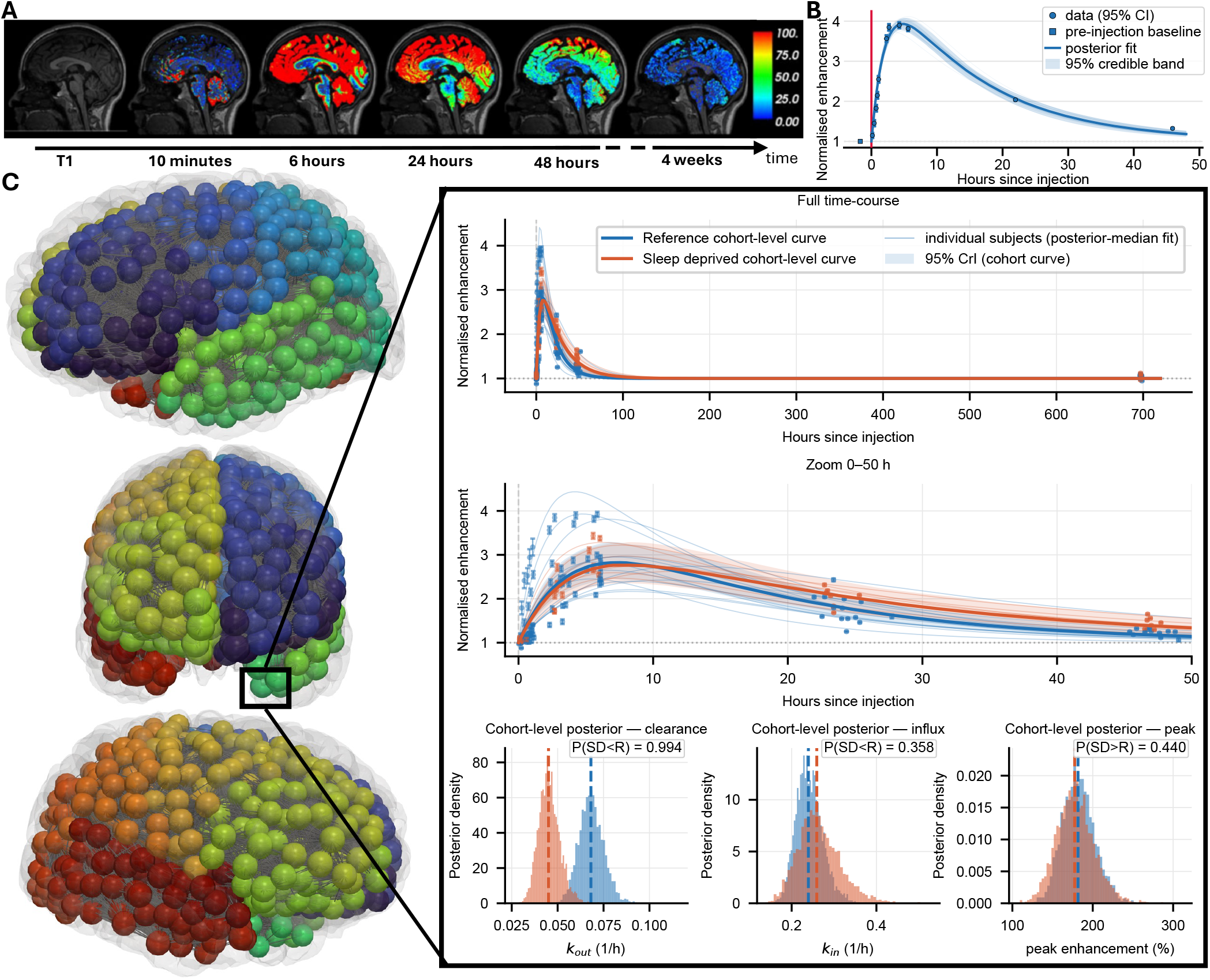
Estimating regional CSF-tracer clearance from contrast-enhanced MRI. (**A**) Sagittal T1-weighted images for a single subject from baseline through four weeks after intrathecal gadobutrol injection, showing tracer enrichment and clearance across the brain (colour bar: percentage T1 signal change). (**B**) The two-compartment fit for one region (entorhinal cortex) in a single subject: baseline-normalised enhancement (points, with 95% confidence intervals from the within-region measurement standard error; square, the pre-injection baseline), 30 individual posterior draws (thin lines), the posterior median fit (solid line) and 95% credible band (shaded), with the injection at *t* = 0 (red vertical line). (**C**) The connectome over which clearance is mapped: 1015 nodes grouped into the 83 regions of interest, positioned in anatomical space and joined by structural connections from diffusion tractography, with the featured region marked by the black box. The insets show that region across the full cohort: each subject’s posterior-median fit (thin lines), and for each cohort the curve implied by its cohort-level parameters, drawn as the median across posterior draws with a 95% credible interval (CrI), over the full time course and zoomed to the first 50 h; and the cohort-level posterior distributions of clearance rate (*k*_out_), enhancement rate (*k*_in_) and peak enhancement for the reference (R) and sleep-deprived (SD) cohorts. Each is annotated with the posterior probability of a difference between cohorts in the direction stated. In the region shown, clearance is lower under sleep deprivation with high probability, whereas enhancement rate and peak enhancement do not differ.

Analysing these clearance maps, we found variation between subjects (Figure 2A–D) and regions (Figure 2E). For the 14 reference cohort subjects, the median clearance rate over regions ranged from 0.031 to 0.065 h^*−*1^, a 2.1-fold difference between fastest- and slowest-clearing subject (Figure 2A). Overall in the reference cohort, the median clearance rate across the 34 cortical regions was 0.049 h^*−*1^, corresponding to a parenchymal tracer half-life of approximately 14 h (interquartile range across regions 0.042 to 0.061 h^*−*1^; 0.051 h^*−*1^ across the 36 Braak-staged regions, Figure 2F). Clearance rates varied 2.5-fold across cortical regions: from slow (0.029 h^*−*1^, half-life 24 h) in the lateral occipital cortex, to fast (0.072 h^*−*1^, half-life 9.6 h) in the medial orbitofrontal cortex, with the entorhinal cortex just behind (0.070 h^*−*1^, half-life 9.8 h). The putamen was clearing slower still, extending the range to 3.3-fold (0.022 h^*−*1^, half-life 32 h).

**Figure 2.**
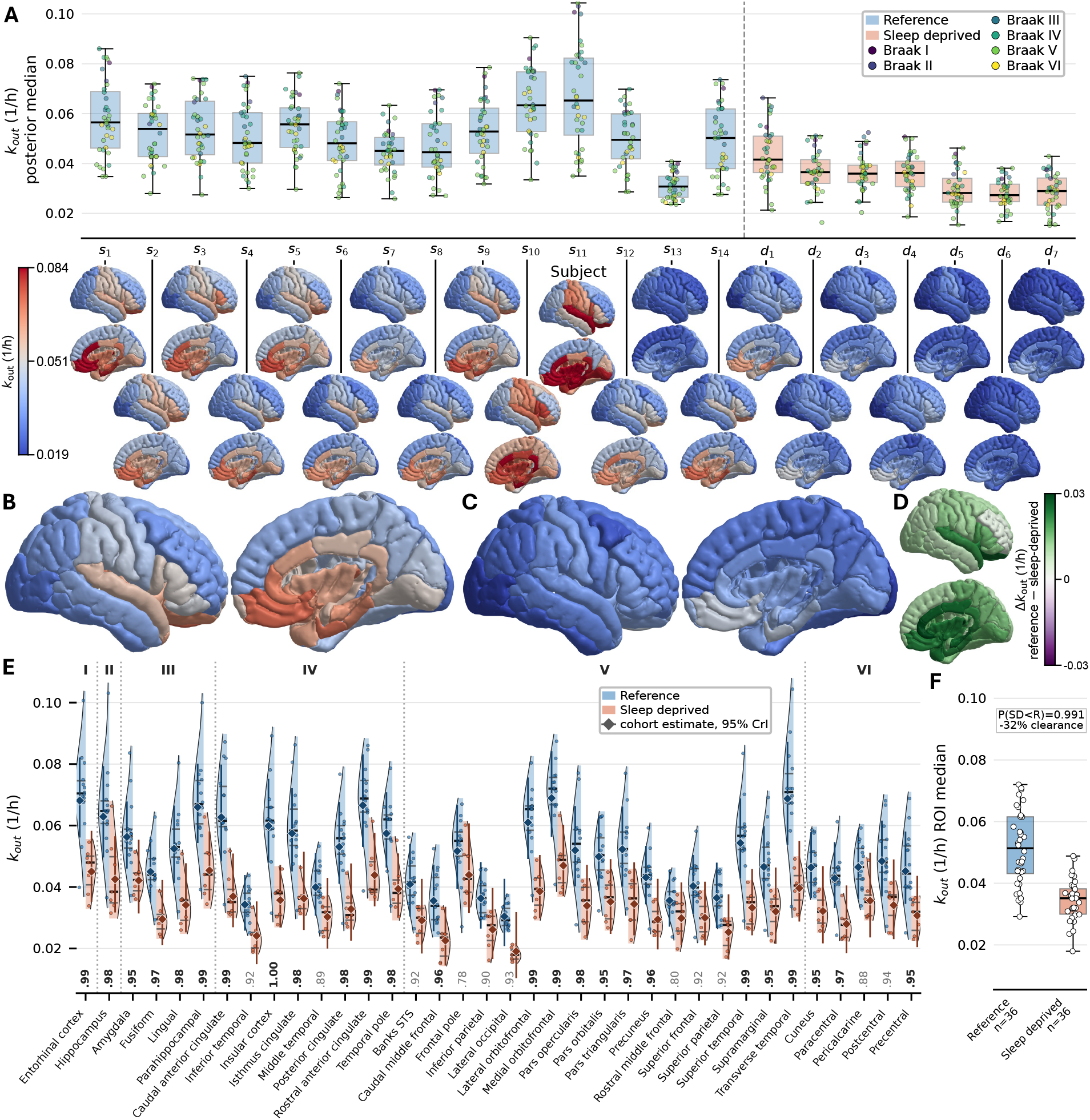
Regional clearance varies over the connectome with consistent spatial patterns across subjects and is reduced after sleep deprivation. (**A**) Per-subject clearance (*k*_out_, posterior median) for the 21 subjects (14 reference *s*_1_–*s*_14_, 7 sleep-deprived *d*_1_–*d*_7_), each box summarising that subject’s rates across the Braak-staged regions, with points coloured by Braak stage, and beneath it that subject’s clearance map on the cortical surface. All maps in A–C share one colour scale, shown in A, on which white marks the reference cohort’s median regional clearance rate (0.051 h^*−*1^). (**B, C**) Clearance maps for the reference (B) and sleep-deprived (C) cohorts, each region coloured by the median across that cohort’s subjects of its subject-level posterior median. (**D**) Between-cohort difference in those regional medians (reference minus sleep-deprived), on a diverging scale centred at zero; green indicates lower clearance under sleep deprivation. (**E**) Regional clearance for both cohorts across all Braak-staged regions, grouped by Braak stage. Violins show the distribution of subject-level posterior medians, and the diamond with bar gives the cohort-level rate and its 95% credible interval. The value beneath each region is the posterior probability that clearance is lower under sleep deprivation (*P*(SD *<* R)), printed in bold where it reaches 0.95. (**F**) Regional clearance summarised across the brain for each cohort, each point one of the 36 Braak-staged regions (the median line of its violin in E); the between-cohort contrast over these regions has posterior probability *P*(SD *<* R) = 0.991, corresponding to a 32% reduction in clearance.

The spatial pattern defined by this ordering of regions by clearance rates was largely consistent across subjects, and can thus be interpreted as a stable property of the clearance maps. Taking the reference cohort first, each subject’s ordering correlated with the average ordering of the remaining subjects (median leave-one-out Spearman’s rank correlation coefficient *ρ*_*s*_ = 0.93, minimum 0.71), and the median pairwise correlation between subjects was *ρ*_*s*_ = 0.87 (*SI Appendix*, S13; Figure S5). The sleep-deprived cohort was similarly consistent (leave-one-out median *ρ*_*s*_ = 0.88, minimum 0.82; pairwise 0.80). Most of the variation in clearance was between regions rather than between subjects: across the reference cohort, 62% of the variance in log clearance lay between regions, 29% between subjects in overall level, and only 9% in subjects departing from the shared regional pattern (*SI Appendix*, S13; Figure S6). Subjects therefore differ in how fast they clear more than in which regions clear fastest (29% of the variance against 9%), and we accordingly summarise each cohort by its median map (Figure 2B, C).

Hierarchical Bayesian inference gives subject-specific clearance rates together with a cohort-level rate in each region. To evaluate the uncertainty of these estimates, we examine the 95% credible intervals (CrIs) associated with the posterior distribution of the cohort-level rates. Across the cortical regions of the reference cohort, the 95% CrI on a region’s cohort-level rate spanned a median 1.64-fold range from its lower to its upper bound (Figure 2E), narrowest in the entorhinal cortex (1.46-fold) and widest in the lateral occipital cortex (2.05-fold). Thus, the entorhinal cortex is the most precisely estimated region in the cohort. Conversely, uncertainty was concentrated in the deep grey nuclei and the lateral occipital cortex. These regions are also those with the lowest tracer enrichment and that show less decay within the observation window, and thus the estimates rely more on partial pooling (*SI Appendix*, S7). We note that the CrIs were wider in the sleep-deprived cohort, with a median range of 1.98-fold.

### Clearance is rapid in the entorhinal cortex and declines across the Braak stage hierarchy

We noted that the entorhinal cortex, a typical site of earliest neurofibrillary change, was among the fastest-clearing of the 42 regions measured. We therefore next examined the distribution of clearance rates specifically in regions implicated in AD: in the entorhinal cortex and in Braak stages I–VI (Figure 3). The reference cohort-level clearance rate (*k*_out_ = 0.068 h^*−*1^, 95% CrI 0.056–0.082) in the entorhinal cortex was 41% above the geometric mean of the cohort-level rates across all 42 regions (95% CrI 25–59%; posterior probability *>* 0.999 that the ratio exceeds one). Moreover, the entorhinal cortex ranked among the fastest-clearing regions in every subject, never falling below the 85th percentile of that subject’s own regional distribution across the 42 regions (Figure 2E). Expressed as the ratio of entorhinal clearance to each subject’s own mean across those regions, which is invariant to overall clearance level, the ratio exceeded one in all 21 subjects (reference median 1.40, interquartile range 1.29–1.43; sleep-deprived median 1.32) (Figure 3A).

**Figure 3.**
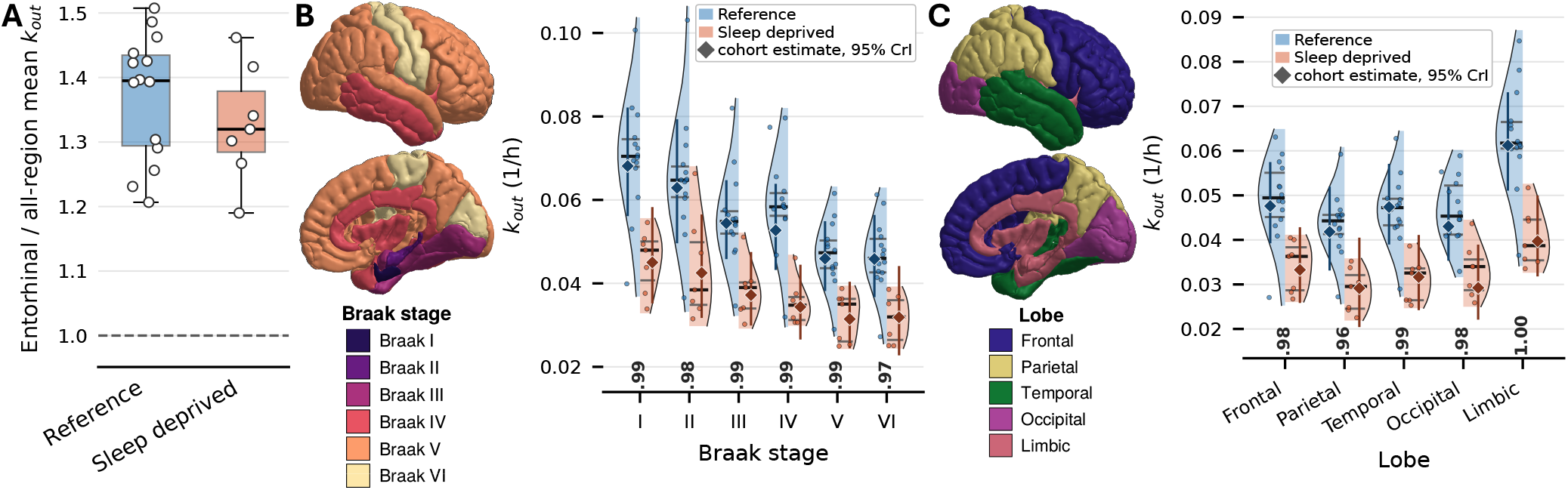
Clearance is rapid in the entorhinal cortex and declines across the Braak stage hierarchy. (**A**) Entorhinal clearance in each subject, expressed as the ratio to that subject’s own mean across all 42 estimated regions. The dashed line at unity marks parity with that mean, which entorhinal cortex exceeded in all 21 subjects. (**B**) Clearance (*k*_out_) aggregated by Braak stage (I–VI; anatomical key at left) for the reference and sleep-deprived cohorts. Each point is the median of one subject’s clearance rates across the regions of that stage; violins show the distribution across subjects and horizontal bars the group median and quartiles. Diamonds with vertical bars show the cohort-level clearance rate for the stage and its 95% credible interval. The value beneath each stage is the posterior probability that clearance is lower under sleep deprivation. (**C**) Clearance aggregated by cortical lobe (anatomical key at left), with cohorts, violins and annotations as in (**B**).

Examining the clearance rates aggregated by Braak stages I–VI, clearance declined overall across this hierarchy (Figure 3B). The rank correlation between a Braak stage’s clearance rate (median across subjects of their stage medians) and its Braak number was *ρ*_*s*_ = *−*0.94, and negative (i.e., declining with stage) in all 14 reference subjects computed within their own maps (median *ρ*_*s*_ = *−*0.89). For instance, stage I cleared 49% faster than stage VI (0.068 versus 0.046 h^*−*1^; 95% CrI +19 to +86%; posterior probability *>* 0.999). Most of the decline occurred at the earliest stages: clearance averaged across stages I–III was 22% above that across stages IV–VI (95% CrI 10–34%; posterior probability *>* 0.999; higher in all 14 reference subjects), and the contrast widened to 35% when only stages I and II were compared with stages III–VI (95% CrI 16–57%; posterior probability *>* 0.999). Adjacent later stages were not separable from one another (posterior probability 0.52 that stage V exceeds stage VI). We also noted that the clearance rates across lobes reflected these features (Figure 3C), with the limbic lobe fastest of the cortical lobes as expected, since it contains the early-Braak medial temporal regions.

### Sleep deprivation reduces clearance without reorganising its spatial pattern

Comparing the reference and sleep-deprived cohorts, clearance was reduced brain-wide after sleep deprivation. Aggregating the 42 regional fits, the sleep-deprived cohort-level clearance was 31% lower than the reference (95% CrI 8–48% lower, posterior probability *P*(SD *<* R) = 0.990), lengthening the half-life implied by the cohort-level rate from 14.4 to 20.8h (Figure 2B, C, D, F). The reduction was widespread: clearance was lower under sleep deprivation in every Braak stage (Figure 3B) and in every cortical lobe (Figure 3C), and in all 42 regions (Figure 2E). Mapped on the cortical surface, the between-cohort difference (reference minus sleep-deprived regional medians) was positive across the entire cortex (Figure 2D).

The tracer enhancement rates *k*_in_ showed no evidence of between-cohort differences (*P*(SD *<* R) = 0.45; cohort-level *k*_in_ +1.5% higher in the sleep-deprived cohort, 95% CrI *−*23 to +36%), and so the reduction in clearance does not appear to be a consequence of changes in influx to the brain. The estimated peak enhancement was instead higher in the sleep-deprived cohort, in 40 of the 42 regions and by 18% at the cohort level (76% above baseline against 64% in the reference cohort; *P*(SD *>* R) = 0.87), and was reached later (16.3h versus 14.2h; posterior probability 0.94). These observations, that sleep-deprived brains showed more enhancement that persisted for longer, are compatible with slower removal during the enhancement phase (*SI Appendix*, S4).

The spatial clearance pattern was preserved after sleep deprivation. Clearance declined overall across Braak stages in both cohorts, with the sleep-deprived profile shifted downward (Figure 3B). The ordering of individual regions largely matched between the two clearance maps (Figure 2B–C; rank correlation *ρ*_*s*_ = +0.85 across the 42 regions). The same differences between the early (I–III) and late (IV–VI) Braak stages were observed in both cohorts: at cohort-level, clearance across stages I–III exceeded stages IV–VI by 22% in both the reference and sleep-deprived cohort (95% CrI 10–34% and 7–40% respectively), and the early-over-late ordering held within every one of the 21 subjects. The size of the reduction varied between 30% and 35% across Braak stages (posterior probability of reduced clearance ≥0.98 in each) without a consistent trend. Across cortical lobes, the cohort-level ordering was identical in the two cohorts (limbic, frontal, temporal, occipital, parietal) and every lobe showed reduced clearance, by 30–35% (Figure 3C). Sleep deprivation thus lowered clearance without reorganising its spatial pattern.

### Subject-specific clearance maps modulate simulated τ propagation across the connectome

Turning from analysis of the clearance rates themselves, we next asked what these clearance differences could imply for the long-term spread of neurotoxic proteins. In particular, we consider simulated propagation of τ proteins, using subject-specific CSF tracer clearance maps as a proxy for τ clearance. After rescaling their timescale as described below, each subject’s clearance map sets the initial regional clearance rates, and the protein concentration is initialised with a localised seed in the entorhinal cortex. We apply a previously published mechanistic model^26^ (Methods) for the coupled dynamics of a dimensionless misfolded protein concentration *p*_*i*_(*t*) and a clearance rate *λ*_*i*_(*t*) (yr^*−*1^) at each node *i* = 1, …, *N* of a connectome *G* :

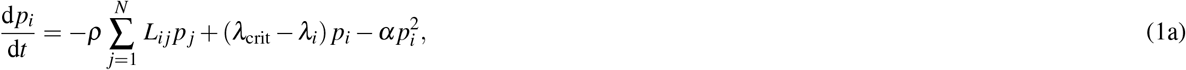

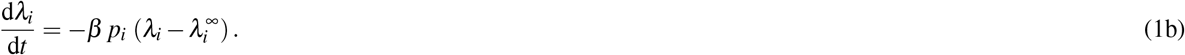

The spreading coefficient *ρ* = 0.01, saturation coefficient *α* = 2.1, clearance-degradation rate *β* = 1 and critical clearance *λ*_crit_ = 0.72, all in yr^*−*1^, were adopted from previous studies^26,27^ (*SI Appendix*, S16, Table S7). The first term on the right hand side of (1a) moves τ protein between connected regions along the fibre tracts of the connectome, at a rate proportional to the concentration difference between them, with 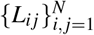 the graph Laplacian of the connectome (Methods), consistent with the prion-like spreading hypothesis for τ^31^. The second term governs local protein accumulation: *λ*_crit_ is the clearance rate at which removal exactly balances aggregation^27^, so when a region’s clearance *λ*_*i*_ falls below it protein accumulates. The third term, 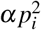, enforces saturation, limiting protein concentration to a maximum level. (1b) models a reduction in the local clearance rate in the presence of misfolded proteins, with 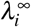 representing the basal clearance rate sustained under the maximal protein concentration.

Simulations were run on a brain network of *N* = 1015 nodes, the finest scale of the Lausanne multiresolution parcellation^32^, with edges derived from diffusion-tractography connectomes^33,34^ (Figure 1C, Methods). Each node belongs to one of the 83 regions at the coarsest Lausanne scale, which are the regions on which the clearance maps are defined. As the CSF tracer clears on a timescale of hours and τ proteins over longer timescales^35^, we rescale each hemispheric clearance rate estimate, before initialising every node with the clearance rate of its parent region. A single scaling constant is applied to every subject in both cohorts, so that all relative differences are preserved. This constant is chosen such that the reference cohort’s mean clearance sits at 0.40 *λ*_crit_, which gives simulated pathology in the reference cohort over a course of several decades, as observed in AD (*SI Appendix*, S16). For each subject, we then simulate disease trajectories as the evolution of the misfolded protein concentration *p* and clearance rate *λ* across the connectome integrated over 100 years and shown to 70 (Figure 4). Simulated fine-scale concentrations are aggregated back to the 83 regions for visualisation. Where a subject’s or a stage’s mean clearance is quoted in this and following sections, it is the mean over the network nodes, so that regions count in proportion to the number of nodes they contain, which approximates their size.

**Figure 4.**
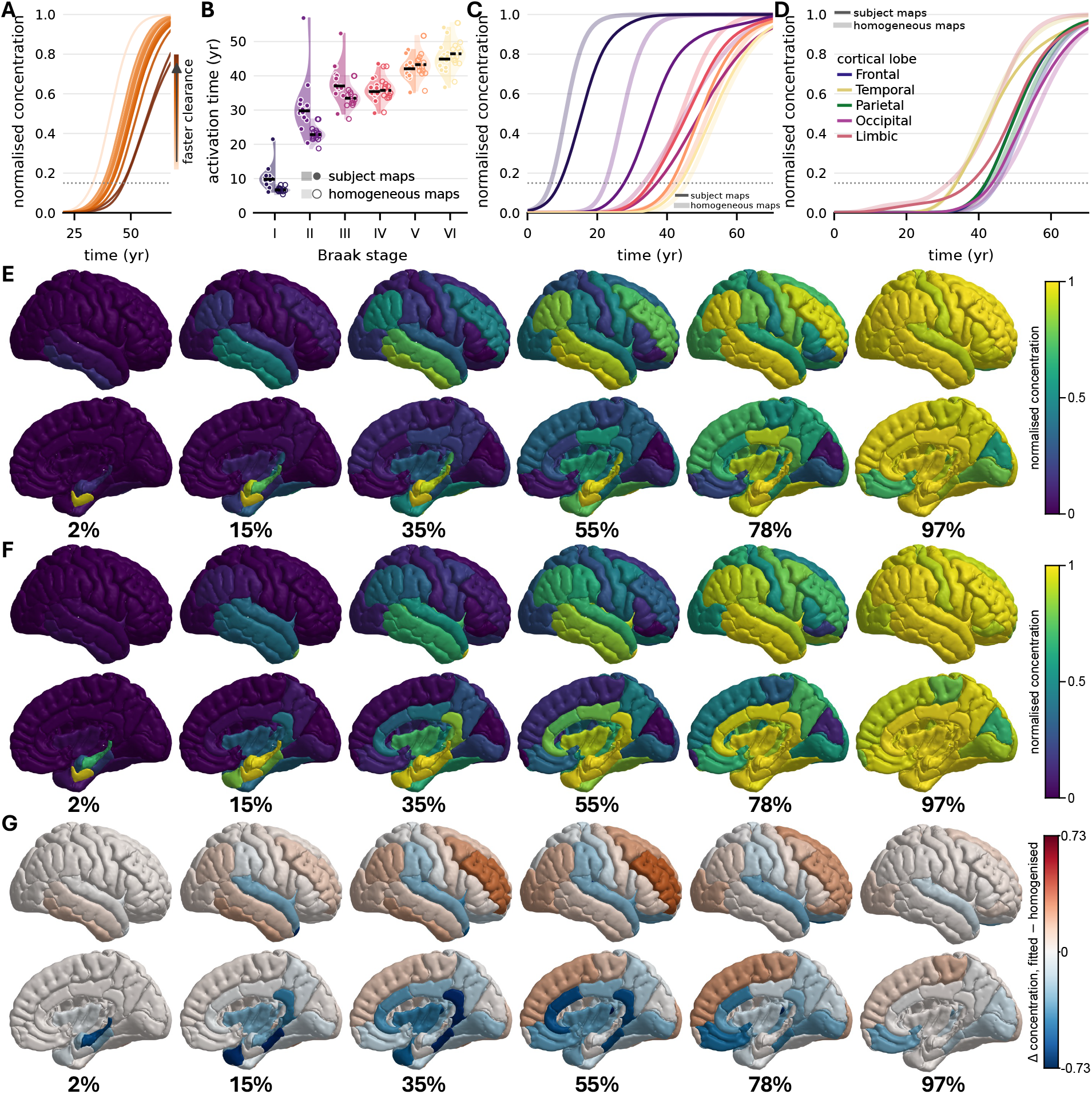
Subject-specific clearance maps modulate simulated τ propagation across the connectome. All panels use the reference cohort (*n* = 14). Each subject is simulated on their own map and the resulting concentration fields are averaged. **(A)** Simulated whole-brain protein concentration over time, one curve per subject, coloured by that subject’s mean clearance *k*_out_, with darker denoting faster clearance. Dotted line, the activation threshold (15%) used throughout. **(B)** Activation time by Braak stage. Split violins: the subject map on the left half, the same map homogenised at matched mean clearance on the right; bars are medians, solid for original and dashed for homogenised. Mean clearance is identical in the two, so the gap between the halves is attributable to spatial pattern alone. **(C)** Mean concentration by Braak stage, averaged across subjects: original subject maps (solid) against the homogenised maps (pale). **(D)** As (C), grouped by cortical lobe. **(E)** Simulated concentration rendered on the cortical surface using the original subject, spatially heterogeneous, clearance maps. Lateral (upper) and medial (lower) views; columns are progression fractions, labelled beneath. **(F)** As (E), with each subject’s map homogenised at their own mean clearance, so mean clearance is held fixed and only the spatial variation is removed. **(G)** The difference between them, original minus homogenised, at matched progression fractions. Red indicates regions where the subject’s own clearance map accumulates more than the homogenised map, blue where it accumulates less. Columns in (E), (F) and (G) are matched on progression fraction rather than on time: each subject is sampled at the time they reach a given fraction of their own final load and the fields are then averaged.

The simulated protein concentrations rose sigmoidally in every subject, with the subject trajectories separated in time by their different initial clearance rates (Figure 4A). To quantify, we examined Braak stage activation times, defined as the time when the simulated mean protein concentration first reached 15% of the model’s asymptotic single-node capacity, a concept analogous to threshold-based definitions of tau positron-emission tomography positivity. The separation in time was ordered by clearance level: the 2.0-fold range in mean *k*_out_ across the 14 subjects in the reference cohort corresponded to a 17.5-year spread in activation times for Braak VI (Figure 4B), the two being correlated at *r* = 0.98. Generally, Braak stage activation times in the reference cohort followed the Braak ordering with one exception (Braak IV), rising from a median of 9.8 yr at Braak I to 44.9 yr at Braak VI (Figure 4B). Between-subject spread was widest at Braak II (36.5 yr) and narrowest at Braak V (12.5 yr), mirroring the coefficient of variation of the stage-mean clearance itself (22.2% and 14.8%; *r* = +0.76 across the six stages). Hence, the stages whose measured clearance varied most between subjects were also those where the activation times vary most.

### The spatial pattern of clearance shapes simulated τ propagation and suppresses initiation

The previous results show that the clearance level, i.e., the mean clearance rate, modulates the simulated τ propagation. Another aspect is how the spatial pattern, as consistently identified in the clearance maps, affects simulation trajectories. For example, in the reference cohort, the entorhinal cortex clears at 1.41 times its subject’s own mean, corresponding to a 27% slower initial protein accumulation than in an average region. To isolate the contribution of the spatial pattern, we first constructed a group of virtual subjects equipped with homogeneous clearance maps; for each subject, we defined the corresponding homogenised clearance map by assigning every node the subject’s mean clearance rate. We then ran the same simulations for the virtual subjects with the homogenised clearance maps as the initial clearance rates, and compared the groups.

In the homogenised virtual reference cohort, we found that Braak I was activated 3.20 yr earlier (interquartile range −3.59 to −2.60) and Braak VI 1.93 yr later (interquartile range +1.56 to +2.33), widening the Braak I to VI interval by 5.24 yr across the 14 subjects (Figure 4B). As such, the spatial pattern compressed the simulated disease course. The pattern also perturbed the staging sequence (Figure 4B, C). Starting from homogeneous clearance rates, the six stages activate in exact Braak order (*ρ*_*s*_ = 1.000), indicating that the connectome and seed alone are sufficient to reproduce the coarse staging sequence, whereas the original subject-specific clearance maps shift Braak IV ahead of Braak III (*ρ*_*s*_ = 0.943). These effects are natural given the decline of clearance rates across Braak stages (Figure 3B): the Braak stage-mean clearance declines from 0.070 to 0.045 h^*−*1^ between Braak I and V, so homogenising reduces protection from early Braak stages and raises clearance across the later Braak stages. At the finer scale, clearance also contributes to which regions within a stage are reached first. For Braak III–VI, faster-clearing regions activate later than other regions in the same stage, with correlations between regional clearance rate and activation time ranging from +0.53 to +0.86 (Spearman, pooled across subjects). Thus, while network topology strongly determined the sequence of simulated progression, regional clearance rates contributed to variation in timing at a finer anatomical scale.

Removing spatial variation in initial clearance also changed where the simulated protein accumulated (Figure 4C–F). The difference between the original and homogenised maps was greatest during the middle of the simulated time course and was more pronounced between Braak stages than between cortical lobes, as expected of a process seeded in the entorhinal cortex. The original subject-specific clearance maps accumulated more proteins than the homogenised ones across the lateral cortex and less in medial temporal regions (Figure 4G). Homogenisation also reduced between-subject variation in the simulated concentration fields: the across-subject standard deviation of regional concentrations fell by 97%, from 0.105 to 0.003 (Figure 4G, at 55% progression). Thus, while the mean clearance is a strong indicator for the timing of simulated disease progression, the spatial clearance pattern contributed to the localisation of the simulated τ-proteins.

We then asked whether the specific arrangement of clearance rates matters independently of their overall spread. Permuting each subject’s clearance rates across regions preserves the distribution exactly, and changes only which region holds which clearance rate. For each subject, we compared simulated τ propagation with their measured map against that with 1,000 permutations of their own per-hemisphere clearance values (*SI Appendix*, S17; Figure S10). The permutations advance the first appearance of simulated τ by a median of 2.9 yr relative to the measured map, in all 14 subjects; the measured map delayed onset more than 98% of its own permutations at the median subject (range 91–100%). Within the permutations themselves, those that carry a higher clearance rate at the entorhinal cortex delay Braak I relative to the rest, and by more the faster that clearance in seed region is (*r* = 0.93, pooled over subjects). When the two hemispheres of each structure were instead permuted together, preserving the measured left–right symmetry, the median delay was 2.7 yr, again in all 14 subjects, with the measured map delaying onset more than 92% of its permutations at the median subject (range 84–98%; *SI Appendix*, Figure S11). Overall, the spatial pattern of the clearance maps estimated from reference (non-AD) subjects delays initiation more than the great majority of rearrangements of the same rates.

### The reduced clearance associated with sleep deprivation advances simulated τ propagation

Finally, we asked what would follow if the clearance differences measured after one night of sleep deprivation were sustained. To examine this, we first mapped how two properties of a clearance map govern progression: its mean clearance and its regional spread, the coefficient of variation of clearance across regions. Taking the reference cohort’s mean map as a template, we rescaled its mean and stretched or shrank its regional deviations from that mean, producing 208 virtual maps that span both properties while keeping the reference arrangement, and recorded the activation time of Braak VI for each (Figure 5A). The contours of simulated Braak VI onset are steep, so a change in mean clearance moves activation times far more than a change in regional spread; and sleep deprivation displaces the reference cohort map almost entirely along the mean-clearance axis.

**Figure 5.**
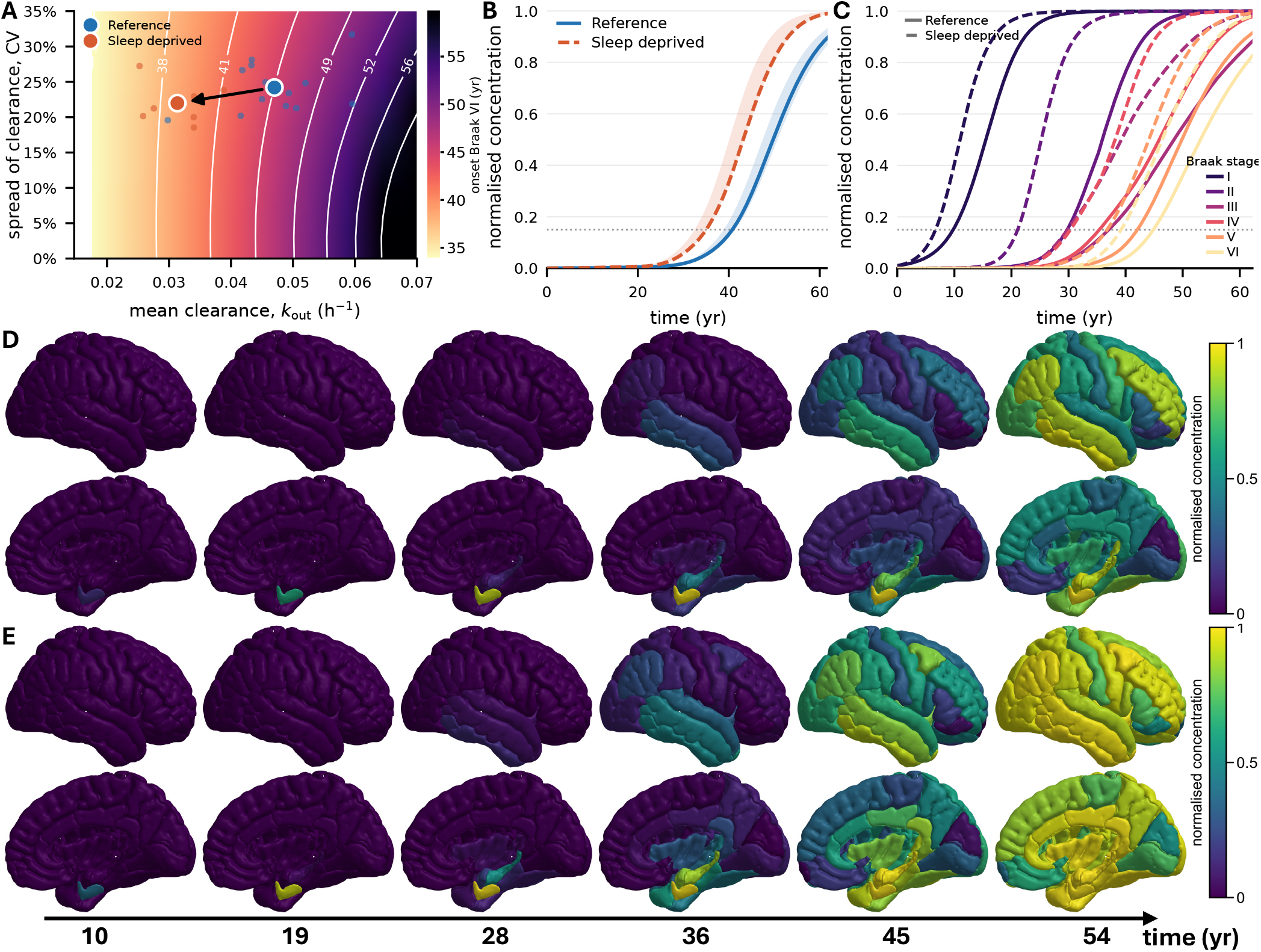
The reduced clearance associated with sleep deprivation advances simulated τ propagation. Reference cohort *n* = 14, sleep-deprived *n* = 7; solid lines and (D) the reference, dashed lines and (E) sleep-deprived throughout. The model is nonlinear in the clearance map, so each subject is simulated on their own map and the resulting fields are averaged. **(A)** Simulated Braak VI onset over the plane of mean clearance and regional spread. The horizontal axis is a map’s mean clearance, in the units of the measured maps; the vertical axis is its coefficient of variation (CV) of clearance across the network’s nodes, with a uniform map at 0%. Small points are individual subjects, large points the cohort means (reference 0.047 h^*−*1^, CV 24%; sleep-deprived 0.031 h^*−*1^, CV 22%), and the arrow the displacement between them. **(B)** Whole-brain normalised concentration over time, cohort medians with interquartile bands. **(C)** The same by Braak stage, cohort medians; colour denotes stage and line style cohort. Dotted lines in (B) and (C) mark the activation threshold (15%) used throughout. **(D)** Simulated concentrations rendered on the cortical surface for the reference cohort, lateral (upper) and medial (lower) views. **(E)** The same for the sleep-deprived cohort. Both surface rows are read at the same six calendar times, 10, 19, 28, 36, 45 and 54 yr, spanning the reference cohort’s median Braak I to latest Braak VI activation times. Colour is normalised concentration on a scale from 0 to 1.

We then simulated τ propagation specifically for each of the subjects in the sleep-deprived cohort, again using the estimated clearance rates as initial conditions to the model (1), and compared the simulated trajectories with the reference cohort. In the sleep-deprived cohort, every Braak stage was activated 3.1–8.7 yr earlier than in the reference cohort simulations (Figure 5B–E). To see whether the advance comes from the lower mean clearance (33%) or the narrower regional spread (9%) of the sleep-deprived maps, we applied each change in turn to every reference subject’s own map, leaving the arrangement of fast and slow regions as measured. Lowering mean clearance alone, with each subject’s regional spread unchanged, advanced every stage by 3.1–7.8 yr (median across subjects), against the 3.1–8.7 yr observed between the cohorts, and accounted for 96–101% of the advance obtained when both changes were applied together. By contrast, reducing the regional spread alone moved any stage by at most 0.8 yr. Sleep deprivation therefore acts on simulated progression through the clearance level, and the loss of protection at the seed follows from that level change rather than from any reorganisation of the map.

## Discussion

Brain clearance is a natural defence against abnormal accumulations of neurotoxic aggregates, and an essential modulating factor of neurodegeneration^36,37^. Glymphatic – extracellular and extravascular – clearance is one of the routes by which solutes leave the brain, and impaired glymphatic function is associated with reduced clearance of Aβ and τ^6,8–11,38^. Here, from consecutive CSF tracer-enhanced MRI^24,39^, we estimate CSF tracer clearance rates in every region of a connectome parcellation in each of 21 subjects. We find that clearance is spatially patterned across the living human brain and largely consistent between subjects. This pattern is distinguished by relatively rapid clearance in the entorhinal cortex and a decline across the stages of the Braak hierarchy. Subjects nevertheless differ in their overall clearance level, and one night of sleep deprivation reduces this level across the brain while largely preserving the spatial pattern. Coupling the clearance maps to a computational model of subject-specific τ protein propagation reveals a distinction between these two properties. The overall level of clearance determines the timescale of simulated pathology, whereas its spatial pattern modifies local vulnerability and the course of progression. Thus, clearance emerges as a spatial property of the brain distinct but complementary to network architecture.

The regional and inter-regional variation in clearance rates estimated and analysed here is broadly consistent with previous human tracer studies. Regional variation in tracer enrichment has been reported in these subjects previously^24^ and in independent cohorts receiving intrathecal gadolinium, where several-fold differences in cortical kinetics have been observed^40^. Spatial variation is also expected and reported for several features proposed to modulate glymphatic transport^6^, including arterial pulsations, cerebral perfusion^41^, glial cell population^42^ and perivascular spaces^43^. At the population level, pharmacokinetic modelling of the same tracer has identified substantial inter-individual variability in CSF-to-blood transport^44^. That analysis and the present work describe different parts of the clearance pathway, but together suggest that inter-individual differences in clearance are not confined to a single compartment. The tracer kinetics themselves are also compatible with previous intrathecal gadolinium measurements. The magnitude and timing of enhancement estimated fall within the range reported by quantitative *T*_1_ mapping and related imaging studies^14,44,45^.

The effects of sleep and sleep deprivation on molecular enrichment and clearance of the brain remain under active debate^21,24,46,47^. We find that after one night of sleep deprivation, clearance is reduced throughout the brain by approximately one third, including at every Braak stage and in every cortical lobe, consistent with previous analyses showing impaired molecular clearance after sleep deprivation^24,39^. An informative contrast emerges from comparison with the original analysis of these data, in which sleep deprivation was not found to measurably alter tracer clearance from the CSF^48^. If both observations hold, this would suggest that the effect of sleep deprivation occurs upstream of net craniospinal CSF clearance, potentially involving transport within the brain parenchyma or local exchange and recirculation between the parenchymal ISF and CSF compartments. This interpretation is consistent with the framework proposed by Wåhlin et al.^49^, in which local CSF–ISF exchange and recirculation may operate partly independently of net craniospinal CSF outflow. We report this as a hypothesis because the studies sample different compartments and use different measurement approaches.

In this study, what is measured is the clearance of an intrathecally injected CSF tracer from each brain region, expressed as a rate, thus quantifying the spatial structure of clearance across the connectome, its consistency across individuals and its reduction after sleep deprivation by comparison between cohorts. What is assumed is that this rate is informative about the regional efficiency with which τ aggregates are cleared, resting on the view that both leave the parenchyma by the extracellular route. What follows from that construction is model-derived: the impacts of clearance rates on the onset and progression of neurodegenerative disease are properties of a simulation and are not direct observations of pathology. Moreover, these simulations should not be interpreted as predicting the consequence of a single night of lost sleep on Alzheimer’s disease progression; they show what the observed reduction in clearance would imply within this simulation framework if it were sustained. Mechanistic modelling provides a plausible basis for this effect, showing that reduced glymphatic flow can impair the removal of larger amyloid-β aggregates^50^. The broader result is that lowering clearance can erode the protection afforded by an otherwise preserved spatial organisation.

Several limitations apply to the tracer measurements and to the rates inferred from them. First, gadobutrol clearance is a proxy for the clearance of neurotoxic proteins rather than a direct measurement of it. The two differ substantially in size (0.6 against 50 kD); while there is evidence of molecular size affecting periarterial enrichment^6^, earlier work suggests transport in the extracellular space is size-independent^51–53^. Second, our estimates require tracer to reach each region: where influx is limited, a low measured rate may reflect poor delivery rather than poor clearance. The least well determined regions are the pallidum and putamen, but none of the principal conclusions rest on them. A regional clearance rate is determined to about ±25% against a threefold spread across regions, and partial pooling attenuates that spread, so the reported regional variation is conservative. The present data do not include regional measures of factors that may influence clearance, such as perfusion, glial density, metabolic rate, or perivascular-space burden, so we cannot determine which of these might explain the observed spatial pattern. Third, the imaging protocol sparsely samples the interval around peak tracer enrichment, making peak timing less precisely resolved than clearance. A Fisher information analysis of the protocol indicates that one additional scan at about 18 h would narrow the interval on cohort-level regional clearance by a third on average, with further scans there adding little. Finally, all subjects were under clinical work-up for suspected CSF disorders [24, Table 1], and their CSF dynamics may therefore not be representative of the general population.

The simulations carry separate limitations. Every subject is simulated on the same group-averaged connectome and from the same entorhinal seed, so between-subject variation arises from clearance by construction. The model also considers a single protein species, whereas Alzheimer’s disease involves interactions between τ, Aβ, vascular processes, and other components that may be cleared through different routes; coupling these mechanisms remains for future work. Finally, the simulations cannot be validated against individual disease trajectories because the subjects do not have longitudinal measures of protein pathology or clinical progression. The results should therefore be interpreted as a mechanistic test of the consequences of the measured clearance maps rather than as subject-specific forecasts.

The broader contribution is a subject-specific estimate of clearance across a connectome parcellation, estimated hierarchically so that each rate carries its own uncertainty, in a form that can parameterise a network model. Structural connectivity is an established determinant of where pathology propagates, while the present results support clearance capacity as a complementary property that may determine how readily different regions permit that pathology to establish and how rapidly the system progresses. This distinction may be relevant to the substantial unexplained variation in the timing of cognitive decline among individuals with preclinical pathology^54–56^. Whether regional clearance provides prognostic information can only be established prospectively, in cohorts with both clearance imaging and longitudinal molecular or clinical outcomes. One testable prediction follows directly from the present results: if rapid entorhinal clearance contributes to protection against the initiation of tauopathy, that regional advantage may be diminished in individuals who subsequently develop Alzheimer’s disease. Because the present maps are derived from subjects without established τ pathology, they describe the clearance architecture present before the classical spatial sequence of tauopathy has emerged.

## Methods

### MRI acquisition and signal intensity interpretation

In reference (*n* = 17) and sleep-deprived (*n* = 7) subject groups, CSF tracer (0.5 mL of 1 mmol/mL, i.e. 0.5 mmol) of the MRI contrast agent Gadovist®, Bayer, Germany (gadobutrol) was injected intrathecally and MR images were collected at multiple time points: at baseline prior to injection, at 0–1.5 h post injection, 1.5–3 h, 4.5–7 h (Day 1), 24 h (Day 2), 48 h (Day 3), and after ≈30 days, as previously reported^24^. During the night between Day 1 and 2 (12–24 h post injection), subjects in the sleep-deprived group were deprived of sleep, while the reference group slept normally. Details of subject demographics are provided in Table S1. The study^24^ was approved by the Regional Committee for Medical and Health Research Ethics (REK) of Health Region South-East, Norway (2015/96), the Institutional Review Board of Oslo University Hospital (2015/1868), the National Medicines Agency (15/04932-7), and was registered in Oslo University Hospital Research Registry (ePhorte 2015/1868). The conduct of the study was governed by ethical standards according to the Declaration of Helsinki of 1975 (and as revised in 1983). Study subjects were included after written and oral informed consent.

### Brain connectome network

The brain’s connectome is organised as a small-world network with high clustering and short path lengths between regions of interest^25,57^. We used a brain connectome network *G* constructed from openly available data^33^ of 426 subjects participating in the Human Connectome project (Figure 1C). The connectomes were constructed using the Lausanne multiresolution atlas^32^, which provides five spatial resolutions, including one with 83 anatomical regions as nodes and one with 1015 nodes nested within those regions^34^. The graph nodes (or vertices), labelled *i* = 1, …, *N*, represent regions of interest (ROIs) in the grey matter. The graph edges represent tractography-derived neuronal fibre tracts between nodes and are weighted by *n*_*i j*_*/l*_*i j*_ where *n*_*i j*_ and *l*_*i j*_ are the average number and length of fibres between regions *i* and *j* respectively^33,58^.

The clearance measurements and rate estimates were defined on the 83-region parcellation. Simulations were run on the high-resolution network consisting of *N* = 1015 nodes, in which each of the 83 regions is subdivided into several nodes. Both graphs are labelled according to the Desikan–Killiany FreeSurfer parcellation^29,30^, so the measured clearance rate for each region is assigned to all of its constituent nodes. This high-resolution graph has been shown to produce more accurate simulations of tauopathy compared to a coarse-grained network with *N* = 83 nodes^34^. For region-based analyses, we also group ROIs into Braak staging regions I–VI using the Alzheimer’s Disease Neuroimaging Initiative (ADNI) mapping of Desikan–Killiany regions to Braak stages.

### Normalised signal intensities and clearance rate maps

To estimate CSF tracer absorption and clearance rates from the brain parenchyma, we use hierarchical Bayesian inference to fit a model to the full contrast-enhancement time course at all post-injection time points. Every available time point was used, including additional acquisitions in the subjects imaged more often than the protocol above. For each subject and each image stack, voxel-wise signal intensities were mapped onto the subject’s Desikan-Killiany parcellation using the median regional intensity value (Figure 1A). Median voxel intensity was used to reduce partial volume effects, though small regions bordering CSF spaces remain susceptible, potentially inflating early signal and steepening the apparent clearance curve. The magnitude of this effect and its consequences for regional comparisons are quantified in the *SI Appendix*, S15; the results reported here were robust in this regard.

To adjust for interscan variations in greyscale, the T1 signal unit at each time point was first normalised by dividing it by the T1 signal of a reference region of interest (ROI) at the same time point. The reference ROI was placed in the posterior part of the orbit, as previously described^24^. These regionally normalised intensities were then divided by the corresponding pre-injection value for that subject and region, yielding, for each subject *s* and region *i*, a baseline-normalised enhancement time course 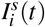 that begins at *I* = 1 and returns toward it as tracer clears (Figure 1B). The 30-day acquisition is not used in the normalisation and so provides an independent check that enhancement does return to baseline (*SI Appendix*, S11).

Subject–region time courses were included on the basis of data availability alone. A subject–region entered the analysis if it had a valid pre-injection baseline, at least three finite time points, and at least one acquisition in each of the 24 and 48 h windows, since the decline between these time points is what identifies the clearance rate. Weakly constrained subject–regions are retained and regularised by partial pooling rather than discarded (*SI Appendix*, S6–S7). Three subjects, all in the reference cohort, had missing acquisitions in the clearance windows and were excluded entirely, leaving 21 of the 24 subjects originally studied (14 reference, 7 sleep-deprived; Table S1). Among these, every subject–region met the criterion, so all 21 retained subjects contribute to every region fit.

To capture the entire time course, including the enhancement phase, we model regional tracer kinetics with a minimal two-compartment model, the analytical solution of a linear system with first-order influx and efflux,

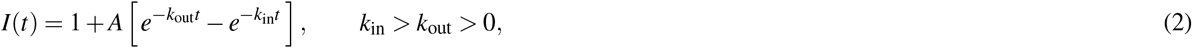

in which *k*_in_ is the effective rate governing tracer enhancement within the region, *k*_out_ is the effective rate governing tracer clearance, and *A* controls the magnitude of enhancement above baseline. The constraint *k*_in_ *> k*_out_ *>* 0 produces an initial rise followed by a slower decay. Compartmental, difference-of-exponentials models of this kind are established for intrathecal gadobutrol kinetics, where a two-compartment model with first-order elimination has been fitted to CSF-to-blood tracer clearance^44,48^. A more in-depth physical interpretation of the model parameters is given in *SI Appendix*, S4.

Clearance rates are estimated using a hierarchical Bayesian model with partial pooling across subjects (Sections S2–S3). Each anatomical structure is fitted three ways: to the left and right hemispheric time courses separately, and to the two pooled into a single bilateral time course. The bilateral fit has its own likelihood and its own pooling and is not the average of the two hemispheric fits. The bilateral fits are used for the regional analyses reported in Results. The hemispheric fits supply the node-level clearance maps for the network simulations, since the connectome distinguishes hemispheres. The two agree closely, the hemispheric mean falling within 0.4% of the bilateral fit at the median across regions (*SI Appendix*, S14). Within a fit, partial pooling operates across all 21 subjects: per-subject log-parameters are Gaussian random effects about a cohort-specific mean, in a non-centred parameterisation, with a single dispersion shared between the reference and sleep-deprived cohorts. No parameters are shared between anatomical regions. The model is sampled in the coordinates (log *k*_out_, log *δ*, log *A*), where *δ* = *k*_in_ *−k*_out_ *>* 0 enforces the rate ordering by construction (*SI Appendix*, S4).

Priors were weakly informative and placed on the log scale. The clearance and time-to-peak priors were informed by published human intrathecal gadobutrol kinetics, and the amplitude prior to the cortical enhancement reported in an independent cohort at the same dose^14,45^; only the observation-noise scale, which is protocol-specific, was set internally. Full specifications, prior-predictive summaries and the sensitivity grid are in the *SI Appendix*, S5. Posteriors were sampled with the No-U-Turn Sampler in NumPyro, four chains of 1,000 warmup and 1,000 retained draws, at identical settings for every region. All fits converged, with no divergent transitions in any region and Monte Carlo error on cohort-level *k*_out_ below 1% of the estimate throughout (*SI Appendix*, S8). Inference was validated by prior-to-posterior contraction, by refitting under thirteen alternative prior, pooling and weighting specifications, by posterior predictive checks, by design-matched parameter recovery, and by simulation-based calibration; the reported results were robust throughout (*SI Appendix*, S7–S12).

The subject-specific posterior median clearance rates yield clearance maps (Figure 2A), displayed on the 83-region parcellation, which were summarised at the cohort level to produce cohort clearance maps for the reference and sleep-deprived cohorts (Figure 2B, C).

### Statistical analysis of the clearance rate maps

Two types of summary are used for different purposes. Descriptive summaries are calculated from subject-level posterior-median rates and reported without intervals. They characterise regional patterns, the spread between subjects, and the consistency of the pattern across subjects. Cohort-level posterior distributions from the hierarchical model are used whenever uncertainty is reported through credible intervals or posterior probabilities.

A subject’s clearance rate in a region is their subject-level posterior median of *k*_out_. Where a subject is summarised across a set of regions (Figure 2A; Figure 3B, C), the summary is the median of these rates across that set. A subject’s overall clearance level, used as the denominator of the within-subject ratio in Figure 3A, is the mean across regions; in the simulations a subject’s level is the mean over the network’s nodes, so that regions count in proportion to the number of nodes they contain. A cohort’s descriptive clearance rate for a region is the median across its subjects of the subject-level medians; this represents the typical subject, is what the cohort clearance maps display (Figure 2B–D), and is the basis of the regional ordering comparisons between cohorts. Where a single number summarises a cohort across regions, it is the median across regions of these regional values. Pattern consistency within a cohort is quantified over the 42 regions as the Spearman rank correlation between subjects’ clearance maps, computed pairwise and, in a leave-one-out form, between each subject and the median clearance map of the remaining subjects; the variance decomposition of log clearance (*SI Appendix*, S13) uses the same subject-level medians.

For each region and cohort, 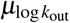 is the mean of the distribution of subject-specific log clearance rates; the between-subject standard deviation is shared across cohorts. When reported as a clearance rate, its posterior distribution is transformed to the *k*_out_ scale to give the cohort-level clearance rate and its credible interval, which quantifies uncertainty in this estimate. Credible intervals are the central 95% posterior quantiles. Their widths are quoted either as a fold-width, the upper bound divided by the lower, or as a percentage either side of the posterior median. Parenchymal half-lives are ln 2*/k*_out_, taken from the same quantity as the rate they accompany. Contrasts in enhancement rate, time to peak and peak enhancement are formed in the same way as those in clearance rate, from the corresponding cohort-level quantities (*SI Appendix*, S4).

We define the between-cohort contrast for a region as 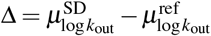 and report it as a percentage change in clearance rate, 100(*e*^Δ^ *−*1), with its posterior median, 95% credible interval, and the posterior probability of reduced clearance under sleep deprivation, *P*(Δ *<* 0 |data), written *P*(SD *<* R) throughout. For a group of regions, such as a Braak stage, a lobe, or the whole set of cortical regions, the cohort-level log rates of the constituent regions are averaged within each posterior draw, a geometric mean on the clearance-rate scale, and contrasts between cohorts or between groups of regions, such as early against late Braak stages, are formed within that same draw. Repeating this across draws yields the posterior distribution of the cohort-level quantity.

### Toxic protein, damage and clearance dynamics

In the network model (1)^26^, two quantities evolve at each node *i*: a dimensionless concentration of misfolded τ *p*_*i*_, which propagates between connected nodes, and a regional clearance rate *λ*_*i*_ (yr^*−*1^), which does not. Clearance removes protein and accumulated protein degrades clearance; *λ*_*i*_ is a single local rate at which protein leaves a region, whatever routes produce that removal. 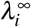 the basal clearance sustained under maximal protein load, is set to zero (numerically 10^*−*6^) as in^26^; *λ*_crit_ is estimated from in vitro aggregation kinetics^27^ . Parameter values and sensitivity are given in *SI Appendix*, S16, Table S7.

To close the system, we impose the following initial conditions:

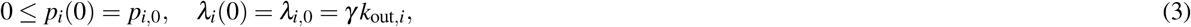

where *k*_out,*i*_ is the subject’s measured clearance rate at node *i* (in h^*−*1^) and *γ* is a single constant that converts it to the model’s units (*SI Appendix*, S16). The epicentre of the misfolded protein population is modelled by setting *p*_*i*_(0) to 1% of the capacity *p*^*∗*^ (defined below) in the entorhinal cortex and zero elsewhere; this seed lies well below the activation threshold, so that Braak I activates by spreading. The graph Laplacian matrix **L** = {*L*_*i j*_ } is used to model diffusion of proteins, from the entorhinal cortex, along axonal pathways on the network and is given by *L*_*i j*_ = *D*_*ii*_ *−A*_*i j*_, where *A*_*i j*_ = *n*_*i j*_*/l*_*i j*_ is the weighted adjacency matrix defined above and 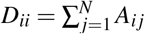 the degree matrix^25,26,34,58,59^.

The first term in (1a) expands to *− ρ*(*D*_*ii*_ *p*_*i*_ *−*∑ _*j*_ *A*_*i j*_ *p* _*j*_), so protein moves between regions in proportion to the difference in their concentrations, weighted by the strength of the connection between them. This represents trans-neuronal spread, in which misfolded protein released by one neuron is taken up by those connected to it^60,61^. The second term is net local growth by prion-like templating: misfolded protein forces normally folded protein into the same abnormal conformation, at a rate proportional to how much is already present^31,62^, offset by clearance acting on the same quantity^5,36^. A region therefore accumulates protein in proportion to how far its clearance falls short of *λ*_crit_; above that rate the term is negative and protein cannot accumulate at all, though every region in our simulations lies below it. The third term is a logistic saturation: a region can sustain only a finite protein load, so accumulation slows and stops at the capacity *p*^*∗*^ (defined below). The second equation (1b) models the feedback of toxic protein on clearance, codifying the observed reduction in clearance in its presence^63–65^.

At any single region, the concentration ultimately approaches 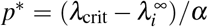, the same for every region, since clearance decays toward 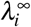 as protein accumulates. For visualisation and comparison between regions, we express protein concentration relative to its asymptotic single-region concentration, *p*_*i*_*/p*^*∗*^, so that 0 represents no misfolded protein and 1 the asymptotic maximum. Activation times are the first times at which the stage-mean concentration reaches 15% of this asymptotic maximum.

### Numerical solution algorithms and software

Clearance rates were estimated in Python using NumPyro, with posteriors sampled by the No-U-Turn Sampler as described in *SI Appendix*, S8. The coupled protein–clearance system (1) was integrated in Python with SciPy’s `solve_ivp` (LSODA), with relative and absolute tolerances of 10^*−*6^ and 10^*−*9^, a maximum step of one two-hundredth of the simulated interval, and output on a grid of 401 points over 100 years. The analysis and simulation code is archived at Zenodo^66^.

### Use of AI tools

Claude (Anthropic; Claude Fable 5.1 and earlier) and ChatGPT (OpenAI; GPT-5-class) assisted with code, checking reported numbers against script outputs, and text editing. All code and values were verified by the authors, who take full responsibility for the content.

## Author contributions

Study conception and design: GSB, TBT, HO, VV, GR, PKE, AG, MER; MRI data collection and analysis: GR, PKE; simulation pipeline: GSB, TBT; analysis and interpretation of results: GSB, TBT, HO, VV, GR, PKE, AG, MER; draft manuscript preparation: GSB, TBT, HO, MER. All authors reviewed the results, edited and approved the final version of the manuscript.

## Competing interests

The authors declare no competing interests.

## Acknowledgments

This manuscript is based on work supported by the EPSRC Centre For Doctoral Training in Industrially Focused Mathematical Modelling (EP/L015803/1) in collaboration with Simula Research Laboratory and the K.G. Jebsen Centre for Brain Fluid Research. M.E. Rognes has received funding from the European Research Council (ERC) under the European Union’s Horizon 2020 research and innovation programme under grant agreement 714892 (Waterscales), and from the Research Council of Norway (RCN) via grants #324239 (EMIx) and #360005 (DigiCells). The work of G.S. Brennan was supported by the EPSRC InFoMM grant EP/L015803/1. The work of T.B. Thompson was partially supported by EP/R020205/1 to A. Goriely and partially supported by the National Science Foundation under grant NSF 2325276 to T.B. Thompson.

## Notes

### Competing Interest Statement

The authors have declared no competing interest.

### Summary of Updates

This version includes substantial updates throughout the manuscript, including revised methodology, updated analyses, and extensive rewriting.

