## Supplementary for "One night of sleep deprivation reduces spatially patterned human brain clearance: implications for simulated tau propagation"

### S1 Subject demographics

Table S1 gives the demographics of the 21 subjects retained in the analysis (Section S6). The source data study is[1].

**Table S1. Demographics of the retained subjects.** Mean ± standard deviation; age range in parentheses; $p$ from the Mann–Whitney $U$ test for age and Body mass index (BMI) and Fisher's exact test for sex. With seven subjects in the sleep-deprived cohort these comparisons are underpowered and are given for description only.

| | Reference | Sleep-deprived | All | $p$ |
|---|---|---|---|---|
| $n$ | 14 | 7 | 21 | — |
| Age, years | 34.7 ± 10.7 (25–60) | 44.7 ± 15.7 (28–65) | 38.0 ± 13.1 (25–65) | 0.11 |
| Female, $n$ (%) | 13 (93) | 6 (86) | 19 (90) | 1.00 |
| BMI, kg/m$^2$ | 29.1 ± 6.6 | 26.2 ± 3.8 | 28.1 ± 5.9 | 0.35 |

### S2 Observation model

For each subject $s$ and region $i$, the baseline-normalised enhancement time course $I_i^s(t)$ is modelled by Eq. [2] (main text). All observations are treated as Gaussian about the predicted curve,

$$I_i^s(t_j) \sim \mathcal{N}\big(\hat{I}_i^s(t_j),\ \sigma_{\text{obs}}\, w_i^s(t_j)\big), \tag{1}$$

where $\sigma_{\text{obs}}$ is a single noise scale learned per region and the weights $w_i^s(t_j)$ set the *relative* precision of each observation. A weight is the standard error of that region's median intensity in that image, $\sqrt{\pi/2}\,\sigma_{\text{vox}}/(\text{median}\sqrt{N_{\text{vox}}})$, where $\sigma_{\text{vox}}$ is the within-parcel voxel standard deviation and $N_{\text{vox}}$ is the number of voxels in the parcel. The normalisation removes the overall scale, which $\sigma_{\text{obs}}$ then estimates from the data. The weights compare observations of the same parcel across time points and subjects, giving more influence to images in which the tracer fills the parcel uniformly. Weights are bounded to $[0.2, 5]$ so that no single observation can dominate a subject's curve or be effectively discarded. The bounds bind for a small minority of observations.

Note that $\sigma_{\text{obs}}$ is not voxel-level measurement error. The weights describe within-image sampling precision (a median of 0.4% of the regional value) whereas the scatter of a measurement about its fitted curve also includes session-to-session registration, segmentation and normalisation variability, and any departure of the true time course from the two-rate form. The fitted $\sigma_{\text{obs}}$ was median 11.7% of the baseline signal across regions (range 4.6–23.9%), lower in the large deep grey parcels and higher in the thin cortical ribbon. We checked the weights against the data. If they capture real differences in precision, then residuals divided by their weight should scatter equally whether the observation was precise or noisy. Splitting the post-injection observations into thirds by weight, the weighted residuals scattered by 1.15, 0.98 and 0.99 in the three groups; without the weights, the same residuals scattered by 0.86, 0.97 and 1.64. The weights therefore capture most of the structure the data show, with one qualification: the third of observations assigned the smallest weights—those the scheme treats as most precise—scatter about 15% more than those weights predict, consistent with a floor from registration and normalisation error that within-image sampling precision cannot see. Removing the weights altogether changes the between-cohort contrast by 3.6% at the median and no conclusion (Section S9).

### S3 Hierarchy

Each region is fitted separately. Within a region, the three subject-level log-parameters are Gaussian random effects,

$$\theta^s = \mu_{c(s)} + \sigma_\theta\, z^s, \qquad z^s \sim \mathcal{N}(0,1), \qquad \theta \in \{\log k_{\text{out}},\ \log\delta,\ \log A\}, \tag{2}$$

with $s$ indexing subjects, $c(s)$ the cohort of subject $s$ ($c \in \{\text{reference}, \text{sleep deprived}\}$), cohort-specific means $\mu_c$, three shared standard deviations $\sigma_\theta$, and $\delta \equiv k_{\text{in}} - k_{\text{out}}$. All are hyperpriors identical across regions, cohorts and subjects, so regional and individual variation enters only through parameters estimated from the data. The non-centred form (a standardised $z^s$ scaled by $\sigma_\theta$) removes the funnel-shaped posterior geometry that hierarchical shrinkage otherwise induces. The two cohorts have separate means but share a single $\sigma_\theta$, so both are shrunk toward their own cohort mean by the same amount. Estimating a separate $\sigma_\theta$ from seven subjects would be poorly determined. A cohort that is more heterogeneous will still produce more widely spread per-subject estimates.

### S4 Parameterisation and physical interpretation of the fitted rates

The three subject-level parameters are sampled on the log scale, $\theta \in \{\log k_{\mathrm{out}},\ \log \delta,\ \log A\}$, and exponentiated to give the rates and amplitude that enter the curve:

$$k_{\mathrm{out}} = e^{\theta_{k_{\mathrm{out}}}}, \qquad \delta = e^{\theta_\delta}, \qquad A = e^{\theta_A}, \qquad k_{\mathrm{in}} = k_{\mathrm{out}} + \delta, \tag{3}$$

so that all three are positive and $k_{\mathrm{in}} > k_{\mathrm{out}}$ by construction. The ordering is an assumption that the data compel: enhancement rises over hours and washes out over days, and the reverse labelling would require tissue to fill over days and empty within hours, contradicting every observed curve and the published time course[2].

The model also implies $t_{\mathrm{peak}}\, k_{\mathrm{out}} < 1$, where $t_{\mathrm{peak}} = \ln(k_{\mathrm{in}}/k_{\mathrm{out}})/(k_{\mathrm{in}} - k_{\mathrm{out}})$: a curve cannot peak later than the reciprocal of its own clearance rate. This held in every posterior draw of every region, even though the pallidum and caudate sample within 5% of the limit in 37% and 30% of their draws respectively, against at most 6% for any cortical region (Section S7).

The fitted curve has a physical reading. If tracer enters a region from the cerebrospinal fluid (CSF) at a rate proportional to the CSF concentration, and leaves at a rate proportional to its own, then

$$\frac{\mathrm{d}T}{\mathrm{d}t} = \kappa C_{\mathrm{csf}}(t) - k_{\mathrm{out}}\, T, \qquad C_{\mathrm{csf}}(t) = C_0\, e^{-k_{\mathrm{in}} t}, \tag{4}$$

where $T$ is the tracer concentration in the region, $C_{\mathrm{csf}}$ that in the CSF supplying it, $C_0$ its value at injection, and $\kappa$ the rate at which the region takes tracer up.

With $T(0) = 0$ the solution to Eq. (4) is

$$T(t) = \frac{\kappa C_0}{k_{\mathrm{in}} - k_{\mathrm{out}}} \left( e^{-k_{\mathrm{out}} t} - e^{-k_{\mathrm{in}} t} \right), \tag{5}$$

which is Eq. [2] (main text)with $A = \kappa C_0/(k_{\mathrm{in}} - k_{\mathrm{out}})$, and it says what the fitted constants mean.

The faster rate, $k_{\mathrm{in}}$, is the rate at which the CSF supply disappears, not a property of the region; the region's own uptake rate $\kappa$ enters only through the amplitude, where it is multiplied by the subject's CSF exposure $C_0$ and divided by the difference of the two rates. The curve peaks at time

$$t_{\mathrm{peak}} = \frac{\ln(k_{\mathrm{in}}/k_{\mathrm{out}})}{k_{\mathrm{in}} - k_{\mathrm{out}}}, \tag{6}$$

with peak enhancement above baseline

$$T_{\mathrm{peak}} = \frac{\kappa C_0}{k_{\mathrm{in}}} \left( \frac{k_{\mathrm{out}}}{k_{\mathrm{in}}} \right)^{k_{\mathrm{out}}/(k_{\mathrm{in}} - k_{\mathrm{out}})} = \frac{A(k_{\mathrm{in}} - k_{\mathrm{out}})}{k_{\mathrm{in}}} \left( \frac{k_{\mathrm{out}}}{k_{\mathrm{in}}} \right)^{k_{\mathrm{out}}/(k_{\mathrm{in}} - k_{\mathrm{out}})}, \tag{7}$$

so the peak enhancement is the uptake-to-supply ratio $\kappa C_0/k_{\mathrm{in}}$ reduced by a factor that depends only on the ratio of the two rates; both $A$ and the peak therefore mix regional uptake with CSF exposure. The clearance rate $k_{\mathrm{out}}$ is the only rate that describes a property of the region alone: it describes how quickly the region empties of tracer, which does not depend on how much tracer reached it, whereas $A$ and $k_{\mathrm{in}}$ do. What $k_{\mathrm{out}}$ does not identify is the route; the loss term in Eq. (4) is the net rate at which tracer leaves the region, not a measurement of any one pathway. Time to peak and peak enhancement are still valid descriptions of the observed curve, and are used as such to compare cohorts and to describe where enhancement is strong or weak; they are not used to infer how well a region takes tracer up, which the curve cannot separate from how much tracer was supplied. Cohort-level enhancement rate, time to peak and peak enhancement are derived within each posterior draw from the cohort-level parameters by $k_{\mathrm{in}} = k_{\mathrm{out}} + \delta$ and Eqs. (6)–(7), and are summarised and contrasted exactly as $k_{\mathrm{out}}$.

The identity $A = \kappa C_0/\delta$ means $A$ and $\delta$ are mechanically linked, whereas the hierarchy gives them independent priors. Simulation confirms the assumption costs nothing at the cohort level: generating synthetic subjects with $A$ derived from $\delta$ rather than drawn independently recovered the same cohort estimates and between-cohort contrast (Section S12).

The measured quantity is tracer-enhanced signal intensity, $I$, not tracer concentration, $T$, but the T1 signal increase is proportional to tracer concentration[1]. Thus $I - 1 = cT$ for a constant $c$ that is absorbed into $\kappa C_0$, so we get the model Eq. [2] (main text); the rates $k_{\mathrm{in}}$ and $k_{\mathrm{out}}$ are unaffected by the scaling.

### S5 Prior specification

Each region's fit has ten hyperparameters: a cohort-specific mean for each of the three subject-level log-parameters ($\mu_{\log k_{\mathrm{out}}}$, $\mu_{\log \delta}$, $\mu_{\log A}$; six in total), one between-subject dispersion per parameter ($\sigma_\theta$, shared by the cohorts; Section S3), and the single observation-noise scale ($\sigma_{\mathrm{obs}}$; Section S2). These take five prior specifications: one for each kinetic parameter, applied identically to the two cohorts' means; one for the three between-subject dispersions; and one for the noise scale. The priors therefore cannot favour one cohort over the other, since the between-cohort contrast compares two means estimated under exactly the same specification.

Every prior is a Normal distribution on the natural-log scale, i.e. log-normal on the scale of the quantity itself, so that rates, amplitudes and scales are positive by construction. All priors are weakly informative, with widths chosen to span the plausible physiological range generously (Table S2). The three kinetic priors are informed by published human intrathecal gadobutrol studies: the clearance rate directly[2,3]; the rate gap $\delta$ through the reported time to peak (Eq. (6)); and the amplitude $A$ through the peak enhancement reported for an independent cohort receiving the same dose[4] (Eq. (7)). The between-subject dispersions $\sigma_\theta$ are deliberately uninformative, with published inter-individual variability fixing only their order of magnitude[3]. Only the observation-noise scale ($\sigma_{\text{obs}}$; Section S2) has no published counterpart in these units, and is anchored instead to this study's own data.

The priors were checked in three ways. First, a prior-predictive check: drawing subject-level parameters from the prior hierarchy and simulating enhancement curves from them, the curves span the plausible range of shapes, with clearance half-lives of 4.7 to 41 h, peak times of 4.4 to 29 h, and peak enhancements of 10% to 280% above baseline (90% ranges), all consistent with published intrathecal gadobutrol kinetics and with the enhancement observed here. With measurement noise added from its own prior, the prior-predictive 90% band contains 92% of the 6,678 post-injection observations, the shortfall lying in the 2–12 h enhancement phase and confined to the deep grey regions, which enhance far less than the cortical regions the amplitude prior was anchored on. Second, prior-to-posterior contraction (Section S7): this shows that the data, not the priors, determine the reported quantities wherever claims are made. Third, prior sensitivity analysis (Section S9): every reported result is preserved under each of thirteen alternative specifications, in which each kinetic prior is widened and shifted in both directions, the dispersion prior is widened, the pooling strength is halved and doubled, and the observation weights are discarded.

The motivation for each prior is summarised below and the values are displayed in Table S2:

***Clearance ($\mu_{\log k_{\text{out}}}$).*** Cortical enhancement falls to near zero by 79 h post-injection, measured by quantitative $T_1$ mapping in a single subject[2]. Approximating the late-time decline by the terminal exponential governed by $k_{\text{out}}$, and interpreting "near zero" as 95 to 99% cleared gives $k_{\text{out}}$ between 0.038 and 0.058 h$^{-1}$, and the prior median of 0.050 h$^{-1}$ (half-life 13.9 h) sits within that range. This is consistent in timescale with population modelling of intrathecal gadobutrol, in which blood concentrations approach zero by 60 h in reference patients[3]. The prior's central 95% range, 0.019–0.133 h$^{-1}$, corresponds to half-lives of 5.2 to 37 h. At the fast bound, this means that tracer is 98% cleared within about 30 h of injection; at the slow bound, just under a quarter still remains at 79 h and clearance takes about nine days. The range is deliberately permissive since prolonged gadolinium presence has been reported after intrathecal administration[5]. Two subjects in our cohort still show incomplete clearance at four weeks (Section S11).

***Time to peak ($\mu_{\log \delta}$).*** The model has no parameter for the time of peak enhancement, but that time follows from the two rates (Eq. (6)), so we specify the prior on the rate gap $\delta = k_{\text{in}} - k_{\text{out}}$ by asking what peak times it implies. Quantitative $T_1$ mapping in a single subject reports cortical grey matter peaking between 10 and 26 h after injection, with white matter peaking later, between 26 and 50 h[2]. The prior lies within the reported cortical window and extends broadly on either side, with a median peak time of 11.5 h, with 90% of its mass between 4.4 and 28.8 h. The breadth is deliberate. Deeper structures are supplied by CSF later than cortex is, and the deep grey nuclei here are fitted with the latest peaks of all regions — the putamen at 39 h, beyond the prior's central range. No scans were acquired during the interval in which cortical enhancement is expected to peak – between 9 and 24 h – so the prior must allow peaks across it. The data determine $\delta$ well in every region (median contraction 0.89, Section S7).

***Peak enhancement ($\mu_{\log A}$).*** The amplitude $A$ is not the peak enhancement itself: the peak is $A$ multiplied by a factor that depends on both rates (Eq. (7)), so a prior on $A$ is best specified by asking what peak enhancement it implies. The prior is taken from the literature: an independent intrathecal gadobutrol cohort receiving the same 0.5 mmol dose reports cortical grey-matter enhancement of 52 ± 54% at its 24 h acquisition[4]. That is a value sampled at a single acquisition, and skips the 9–24 h window in which cortical enhancement is expected to peak[2], so it understates the true peak enhancement. Scaling by 1.2 gives a target peak enhancement of 62%, which is the median peak the prior produces when curves are simulated from all three priors together. The factor is an assumption, but Section S9 shows that shifting the centre by half a log unit in either direction, almost three times its size, leaves every reported result unchanged.

The prior width admits the peak enhancements observed in the present cohort, which runs somewhat higher than the published one: at the corresponding acquisition (24 h), cortical grey-matter enhancement in the reference cohort we fit to is 67 ± 37% ($n = 14$), a 15-point difference, less than half the between-subject spread within either cohort. The prior's 90% range on peak enhancement is 10% to 280%. The highest single enhancement observed in this study (347%, over all subject-regions) sits at the 97th percentile, so even the most extreme enhancement observed retains prior support.

***Between-subject dispersion ($\sigma_\theta$).*** Subjects differ from one another, and $\sigma_\theta$ quantifies how much. Each subject's clearance rate, rate gap and amplitude sit around their cohort's average, and $\sigma_\theta$ is the standard deviation of that scatter on the log scale.

The three parameters share the prior on $\sigma_\theta$. Its median corresponds to a typical subject lying within about 50% above or below their cohort average. The central 95% range of the prior spans between-subject dispersions from about ±25% to a factor of 2. Both cohorts share a single $\sigma_\theta$, so sleep deprivation is modelled as shifting the average rather than the spread.

Individuals differ substantially in intrathecal gadobutrol kinetics: coefficients of variation of 53 to 65% in how fast the tracer is absorbed, when it peaks and how high it reaches[3], which is 0.50 to 0.59 on the log scale. Those measurements describe transfer from CSF to blood rather than regional clearance, so they fix only the order of magnitude for our prior, which is centred at 0.4 with a range wide enough to cover them. The fitted dispersions had cross-region medians of 0.30 for $\log k_{\text{out}}$ (range 0.21–0.48), 0.46 for $\log \delta$ (0.29–0.80) and 0.35 for $\log A$ (0.24–0.69), straddling the prior centre and all below the published range. Halving or doubling that centre leaves the reported results essentially unchanged (Section S9).

***Observation scale ($\sigma_{\text{obs}}$).*** This is the scatter of a regional measurement about the fitted curve, as a fraction of baseline signal. The weights, derived from within-image precision, set the relative reliability of observations; $\sigma_{\text{obs}}$ sets the overall scale (Section S2). The four-week acquisitions give an empirical anchor for that scale: by then tracer has largely cleared, so the measurement should return to baseline. Its scatter about that baseline has a standard deviation of 5.6% once the two subjects whose tracer had not cleared are excluded (Section S11). The prior has median 8%, close to that figure, with a central range of 3 to 21%. The fitted value is higher, at 11.7%, since a measurement scatters more when the tissue is enhanced than it does about a static baseline (residuals about the fitted curve scatter about 25% more, in absolute terms, at high enhancement than at low; Section S10). The posterior predictive checks of Section S10 confirm this fitted scale is calibrated (standardised residual SD 1.05, 95% coverage 96.1%).

**Table S2. Prior specification.** Priors are placed on the natural-log scale of each quantity, which keeps it positive and symmetric in relative terms; implied medians and central ranges are shown back-transformed to natural units. Anchors are identified here and justified in Section S5: the three kinetic priors follow published intrathecal gadobutrol kinetics, while the observation-noise scale, which has no published counterpart in these units, is informed by this study's own data. Cohort means are estimated separately for the two cohorts; all priors are global across regions, cohorts and subjects.

| Quantity (plain meaning) | Anchor | Prior median | Prior range |
|---|---|---|---|
| *Kinetic parameters* | | | |
| Clearance rate $k_{\text{out}}$ (how fast tracer washes out) | near-complete clearance by ∼79 h[2,3] | $0.050\,\text{h}^{-1}$ | $0.019$–$0.133\,\text{h}^{-1}$ (95%) |
| as tissue half-life $t_{1/2} = \ln 2/k_{\text{out}}$ | | 13.9 h | 5.2–36.5 h |
| Time to peak (when enhancement is highest; set through the prior on $\log \delta$) | cortical grey-matter peak 10–26 h, deeper structures later[2] | 11.5 h | 4.4–28.8 h (90%) |
| Peak enhancement (height of the signal rise, as a fraction of baseline; set through the prior on $\log A$) | cortical enhancement in an independent cohort (52%), corrected for single-timepoint sampling[4] | 0.62 | 0.10–2.80 (90%) |
| *Between-subject spread and observation noise* | | | |
| Between-subject spread (how much individuals differ; one prior for all three kinetic parameters) | published inter-individual variability in intrathecal gadobutrol kinetics[3] | 0.40 | 0.22–0.72 (95%) |
| Observation scale $\sigma_{\text{obs}}$ (scatter of a measurement about the fitted curve) | observed four-week scatter about baseline (Section S11) | 8% | 3–21% (95%) |

### S6 Inclusion and fit coverage

Inclusion was evaluated per subject–region from data availability. A subject–region entered the fit if it had a valid pre-injection baseline, at least three finite points, and at least one scan in each of the 24 and 48 h acquisitions (actual acquisition times 22.0–27.7 h and 45.4–50.9 h across subjects) since the decline between these points is what identifies $k_{\text{out}}$. Weakly informative subject–regions are retained and regularised by partial pooling rather than discarded.

Three subjects (all reference cohort) had no scans in the clearance windows and were excluded entirely, leaving 21 subjects (14 reference, 7 sleep-deprived). Among these, every subject–region met the criterion: all 882 bilateral curves (21 subjects ×

42 regions) entered the fits, with no exclusions for a missing baseline, insufficient timepoints, or inadequate sampling of the clearance phase.

### S7 Practical identifiability

Identifiability was quantified at the cohort level by prior-to-posterior contraction, $1-\left(\frac{\sigma_{\text{post}}}{\sigma_{\text{prior}}}\right)^2$, where $\sigma_{\text{post}}$ and $\sigma_{\text{prior}}$ are the posterior and prior standard deviations, respectively. Contraction of the cohort-level clearance rate was high throughout: median 0.934 across the 42 regions, minimum 0.594 (pallidum). The rate gap $\delta = k_{\text{in}} - k_{\text{out}}$ contracted almost as strongly (median 0.890, minimum 0.639). The amplitude $A$ contracted less (median 0.779, minimum 0.246), falling below 0.5 in the five deep grey structures with least signal and in lateral occipital cortex, as expected of a parameter that combines uptake, exposure and the rate gap in regions with least signal (Section S4). No principal clearance conclusion depends on the amplitude estimate, and in the entorhinal cortex all three parameters contract above 0.93. The weak-identification criteria used here were contraction of the clearance rate below 0.5, or a posterior correlation between $\log k_{\text{out}}$ and $\log \delta$ exceeding 0.95 in magnitude; neither was met in any region.

That second criterion concerns how much the two rates, $\log k_{\text{in}}$ and $\log k_{\text{out}}$, constrain one another. Within a single enhancement curve a slower clearance can be partly mimicked by a faster influx, so the two are not separately determined: their posterior correlation has median $-0.52$ across regions. The reported credible intervals for $k_{\text{out}}$ are marginal posteriors and so already integrate over this uncertainty in the enhancement rate.

The least well identified regions are the deep grey nuclei, which occupy three of the four lowest contraction values (Table S3): pallidum (0.59), putamen (0.74) and caudate (0.77). These structures lie furthest from the CSF surfaces that supply them, so they enhance late and weakly: the putamen's fitted peak enhancement is 9% above baseline, against an observation scatter of 5%, where entorhinal cortex reaches 182% against 13%. Contraction tracks this signal-to-noise ratio (the posterior median of the cohort-level peak enhancement divided by that of the observation scale) across all 42 regions ($r = 0.87$; Figure S1, left). With so little signal, the sparse sampling that constrains the cortical curves leaves the deep grey curves underdetermined, and their fitted peak times approach the upper limit the two-rate model allows, $t_{\text{peak}} < 1/k_{\text{out}}$ (Section S4): in more than a third of posterior draws for the pallidum, and nearly a third for the caudate, the peak lies within 5% of that limit. The least well identified cortical structure is the lateral occipital cortex, which has the weakest enhancement of any cortical region (19% above baseline; Figure S1).

The clearance estimates for these weakly enhanced regions are therefore drawn more toward the prior. For the pallidum, its contraction value implies a pull of +22%; refitting with a clearance prior shifted by half a log unit moves it by +26% and −19%, the largest response of any region (Section S9); and simulation at the study's own design recovers the deep low-signal regime [+11% to +36%] high (Section S12). Lateral occipital cortex is treated the same way: its fits fail to converge under two of the thirteen prior variants, and where they converge its cohort contrast is the most prior-dependent of any region, moving by up to 0.33 in log units when the clearance prior is widened, against at most 0.10 for any other region (Section S9). No conclusion rests on an estimate from these regions. The five deep grey structures lie outside the Braak-staged set, enter the whole-brain contrast as five regions of 42, and are named in the main text only to mark the extremes of the clearance range. Lateral occipital cortex is one of 17 regions in Braak stage V and one of four in the occipital lobe; its cohort-level rate is shown with its credible interval (Figure 2E).

Partial pooling draws each subject toward its own region's cohort mean, so the ordering of regions by clearance is unaffected and the reported regional spread is if anything conservative. The regions it most affects are the slow, low-signal deep grey structures, in which simulation shows clearance biased upward toward the prior (Section S12). The entorhinal elevation on which the regional claims rest lies in the best-identified class (contraction 0.96), moves by less than 2% under every prior variant, and is recovered in simulation in all 12 replicates with a bias of −2% (Section S12).

### S8 Sampling and convergence

Posteriors were sampled with No-U-Turn Sampler (NUTS) in NumPyro: four chains of 1,000 warmup and 1,000 retained iterations (4,000 draws), target acceptance 0.99, maximum tree depth 10, and chains initialised at the prior median. Every fit used these same settings—the 42 bilateral regions (34 cortical and 7 subcortical structures fitted as left–right pairs, plus the midline brain stem), the whole-cortex aggregate, and the 82 per-hemisphere fits (41 paired regions, left and right) used for the surface maps and simulations, 125 in all—and all converged, with no divergent transitions in any fit. For the cohort-level clearance rate, the worst $\hat{R}$ was 1.003 and the minimum bulk effective sample size 1,996 (median 3,006), giving a Monte Carlo standard error of 0.23% of the estimate at the median and 0.63% at worst, negligible against every effect reported here. Across all ten sampled hyperparameters of all 125 fits, the worst $\hat{R}$ was 1.007 and the smallest effective sample size 689, both occurred for between-subject dispersion parameters in per-hemisphere fits (right caudal anterior cingulate and left superior parietal), which are the slowest-mixing quantities in a hierarchical model and are not themselves reported. No region required longer runs or altered settings.

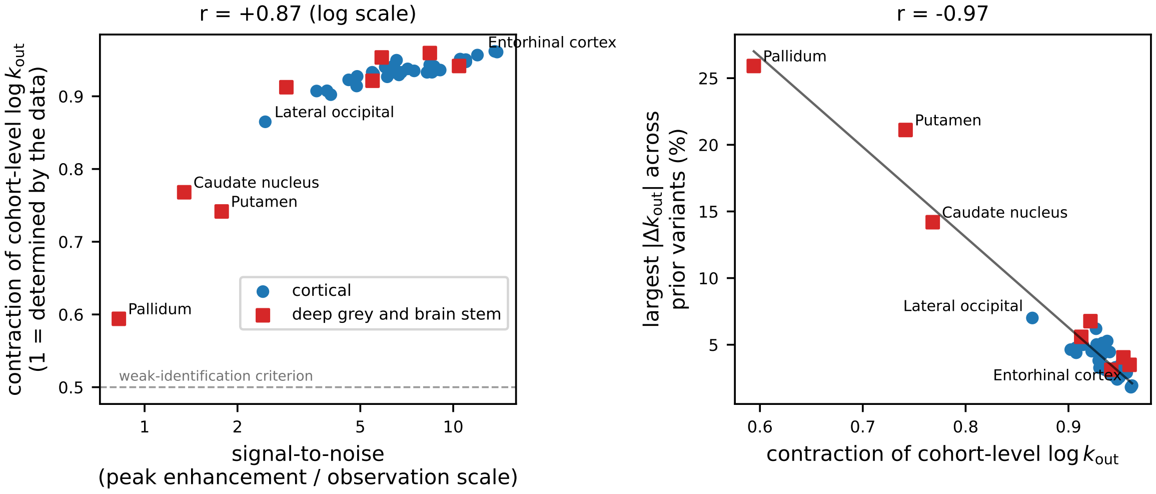


**Figure S1. Identifiability follows signal, and prior dependence follows identifiability.** Left: prior-to-posterior contraction of the cohort-level clearance rate against each region's signal-to-noise ratio, the fitted peak enhancement divided by the fitted observation scale. The deep grey nuclei (squares) have the least signal and the lowest contraction; lateral occipital cortex is the only cortical region among them; no region falls below the weak-identification criterion (dashed). Right: the largest change in each region's cohort-level clearance rate across the ten prior variants of Section S9 (the nine kinetic-prior variants and the widened dispersion hyperprior), against the same contraction; the line is a least-squares fit. Regions the data determine least are the ones a changed prior moves most, and the entorhinal cortex is among the least moved.

**Table S3. Prior-to-posterior contraction of the cohort-level clearance rate (reference cohort)**, showing selected regions from the least to most data-determined. Contraction is $1 - \left(\frac{\sigma_{post}}{\sigma_{prior}}\right)^2$: values near 1 indicate a rate determined by the data, values near 0 one carried by the prior. The final column gives the proportion of posterior draws in which the fitted peak time lies within 5% of the upper limit the two-rate model allows, which separates the three least-identified nuclei from the rest.

| Region | Contraction | Draws near the peak-time limit |
|---|---|---|
| *Least data-determined* | | |
| Pallidum | 0.594 | 37% |
| Putamen | 0.742 | 10% |
| Caudate nucleus | 0.768 | 30% |
| Lateral occipital cortex | 0.865 | 5% |
| All 42 regions (median) | 0.934 | — |
| *Most data-determined* | | |
| Accumbens area | 0.953 | 0% |
| Medial orbitofrontal cortex | 0.957 | 0% |
| Amygdala | 0.959 | 0% |
| Entorhinal cortex | 0.961 | 0% |
| Parahippocampal cortex | 0.962 | 0% |

### S9 Robustness to priors, pooling strength and weighting

Every region was refitted under thirteen alternative specifications, with identical data and random seeds. Each of the three kinetic priors was widened, by doubling its standard deviation, and shifted in each direction, by half a log unit in its centre; the between-subject dispersion hyperprior was widened and its centre halved and doubled; and the observation weights were discarded entirely. Shifting a prior's centre by half a log unit moved the implied clearance half-life from 13.9 to 8.4 and 23 h; the implied peak time from 12.7 to 10.6 and 14.7 h; and the implied peak enhancement from 62% to 38% and 102% above baseline, a factor of 1.65 in each direction, spanning and in places exceeding the range supported by published kinetics.

Every reported result survives every specification (Table S4). The estimated reduction in clearance under sleep deprivation ranges from 29.5% to 35.0% against a baseline of 31.4%, with the direction preserved in every converged fit; the entorhinal

**Table S4. Robustness of the reported results to the prior, pooling and weighting specification.** Each row is a complete refit of every region under one alternative specification, with identical data, weights and random seeds. The first three columns give the resulting change in cohort-level clearance, separated into the 33 cortical regions and the 8 deep grey structures (seven subcortical nuclei and the brain stem), which the identifiability analysis (Section S7) predicts to be the most prior-dependent; the region carrying the largest deep grey shift is named. *Entorhinal elevation* is the elevation of entorhinal clearance above the whole-brain mean, with its rank among the 41 regions in parentheses; *reduction* is the fall in cohort-level clearance under sleep deprivation given by that refit aggregated over the regions as in the main text. Statistics are computed over the 41 regions that converged under every specification, that is all regions except lateral occipital cortex, which failed to converge under two of the thirteen; over all 42 regions the baseline reduction is 31%, as quoted in the main text.

| | median $\lvert\Delta k_{out}\rvert$ | | largest deep grey | | |
|---|---|---|---|---|---|
| Specification | cortical | deep grey | shift (region) | Entorhinal elevation | Reduction |
| Baseline | — | — | — | +35.6% (3) | 31.4% |
| *Clearance prior* | | | | | |
| widened ($\sigma_{prior} \times 2$) | 0.7% | 0.9% | 21.1% (putamen) | +37.3% (3) | 35.0% |
| centre +0.5 | 2.9% | 4.0% | 25.9% (pallidum) | +33.0% (3) | 29.5% |
| centre −0.5 | 3.1% | 4.0% | 19.3% (pallidum) | +38.0% (2) | 34.1% |
| *Rate-gap prior* | | | | | |
| widened ($\sigma_{prior} \times 2$) | 0.7% | 2.2% | 11.3% (pallidum) | +34.2% (3) | 31.3% |
| centre +0.5 | 1.1% | 1.3% | 2.4% (brain stem) | +36.3% (3) | 31.8% |
| centre −0.5 | 1.1% | 1.1% | 2.5% (brain stem) | +35.1% (3) | 31.1% |
| *Amplitude prior* | | | | | |
| widened ($\sigma_{prior} \times 2$) | 2.8% | 4.9% | 14.5% (pallidum) | +35.4% (3) | 32.3% |
| centre +0.5 | 3.0% | 4.4% | 6.8% (brain stem) | +32.8% (3) | 30.0% |
| centre −0.5 | 3.0% | 4.9% | 6.0% (putamen) | +38.4% (2) | 32.9% |
| *Between-subject dispersion* | | | | | |
| hyperprior widened | 1.6% | 1.5% | 14.2% (caudate) | +36.6% (3) | 33.1% |
| pooling strengthened | 2.7% | 2.2% | 6.1% (pallidum) | +33.9% (3) | 34.0% |
| pooling weakened | 2.4% | 2.1% | 10.3% (pallidum) | +37.5% (2) | 30.5% |
| *Observation weights* | | | | | |
| discarded | 6.8% | 5.6% | 13.8% (hippocampus) | +40.2% (3) | 31.9% |
| Range | 0.7–6.8% | 0.9–5.6% | — | +32.8–40.2% | 29.5–35.0% |

margin over the whole-brain mean ranges from +32.8% to +40.2% against +35.6%; the decline of cohort-level clearance across Braak stages is monotone under every prior and pooling specification ($\rho_s = -1.00$ over the six stages), with only the unweighted fit exchanging stages III and IV ($\rho_s = -0.94$); and the regional ordering is essentially unchanged throughout, correlating with the baseline fit at $\rho_s \geq 0.977$. The largest single influence is the observation weights, whose removal shifts cortical clearance rates by 6.8% at the median, more than twice the effect of shifting any prior's centre by half a log unit, while moving each region's between-cohort contrast by only 3.6% at the median and the whole-brain reduction from 31.4% to 31.9% (last column), and still without altering any conclusion.

Prior dependence falls where the identifiability analysis predicts it should: across the 42 regions, the largest shift in a region's cohort-level clearance rate under any prior variant correlates with its contraction at $r = -0.97$ (Section S7; Figure S1, right). Cortical estimates move by 3.1% or less under every prior perturbation, while the deep grey structures move further, and the largest movements are carried by the regions with the lowest contraction: the pallidum under a shifted clearance prior (25.9%) or a widened amplitude prior (14.5%), the putamen under a widened clearance prior (21.1%). The effect is confined to those structures, with median deep grey shifts at or below 5.6%. Lateral occipital cortex, the least well-identified cortical region and the weakest-enhancing, failed to converge under two of the thirteen specifications and is excluded from Table S4 entirely; under weakened pooling its cohort contrast moves by +0.36 in $\Delta \log k_{out}$ (and this is one of the specifications in which it did not converge), indicating that with so little enhancement of its own that region's contrast is carried largely by the cohort structure. No principal conclusion rests on the lateral occipital estimate.

### S10 Posterior predictive checks

Standardised residuals pooled over posterior draws ($1.3 \times 10^6$ draws across the 42 regions, post-injection points only) had standard deviation 1.045 against a target of 1, with per-region values spanning 1.01–1.06: no region is systematically misfitted. Posterior predictive coverage at the nominal 95% level was 96.1%, so marginally conservative (Figure S2, left). Residuals were mildly heavy-tailed (excess kurtosis +0.71, skew +0.35), localised to the low-enhancement regions in which isolated points carry disproportionate weight, and do not distort the reported intervals. The weight-tercile validation reported in Section S2 was computed from the same residuals.

One systematic feature is present, and the acquisition times localise it. The residual mean was −0.22, and negative in all 42 regions, but it is not uniform: observations within 2 h of injection lie 0.72 standard deviations below the fitted curve, those between 2 and 100 h lie on it (−0.02), and the four-week scans lie 0.28 above it, consistent with their median enhancement of 1.031 (Figure S2, middle; Section S11). The two-rate curve therefore rises slightly too steeply in the first two hours and returns to baseline slightly too completely by a month, neither of which is the 24–48 h decline that determines $k_{\mathrm{out}}$. The between-cohort contrast, a difference between cohorts fitted under the identical constraint, is insensitive to a shared offset of this kind.

Seen by fitted value rather than by time (Figure S2, right), the same offset sits at low enhancement and vanishes above about twice baseline, and the residual scatter rises modestly with signal, from an SD of 0.93 in the lower half of fitted values to 1.15 in the upper half. The lowest bin in that panel is the four-week scans, which sit slightly above the curve; the offset below the curve is the early rise.

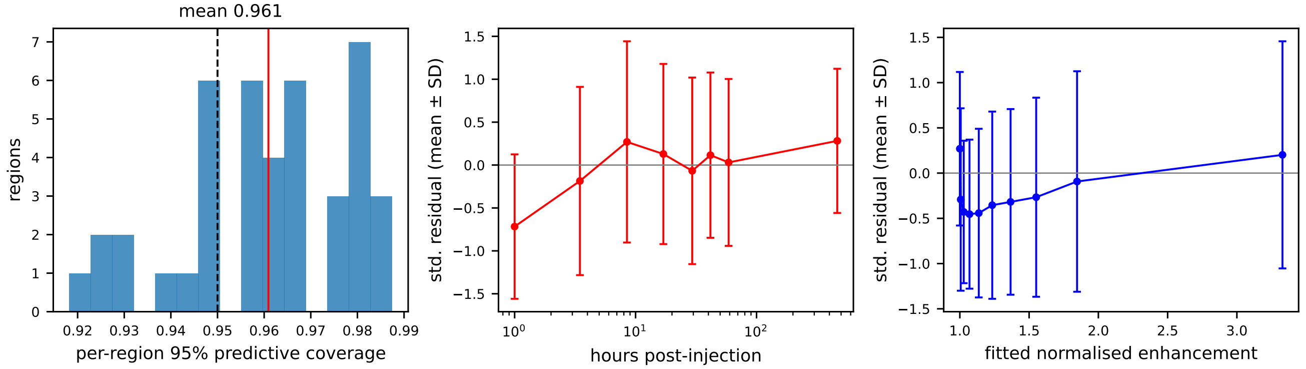


**Figure S2. Posterior predictive checks.** Left: per-region coverage of the 95% posterior predictive interval (dashed, nominal; solid, mean over the 42 regions). Middle: standardised residuals, mean ± SD, by acquisition time; the model overpredicts the early scans and underpredicts the four-week scans, and the 10–60 h window that determines the clearance rate lies on the curve. Right: the same residuals by fitted enhancement, in ten equal-count bins. Post-injection observations only, 200 posterior draws per region.

### S11 Robustness to individual subjects and return to baseline

Because the sleep-deprived cohort is small ($n = 7$), we verified that no individual subject drives the between-cohort contrast. Three regions were chosen for this: the entorhinal cortex, on which the regional claims rest; the hippocampus, the other early-Braak medial temporal structure; and the whole cerebral cortex as a single fit. For each, the model was refitted with every subject removed in turn, 21 refits per region, one for each single-subject deletion (Table S5).

The between-cohort contrast was robust throughout: across all 63 refits the posterior probability of reduced clearance remained above 0.92, and the median contrast ranged from −0.328 to −0.484 in log units (a 28 to 38% reduction) against full-data values of −0.414, −0.389 and −0.392 (34, 32 and 32%) for the entorhinal cortex, hippocampus and cortical grey matter respectively. No omission changed the direction of the effect in any region.

The deletions that weaken the contrast most are from the sleep-deprived cohort. In every region the weakest refit omits a sleep-deprived subject, and in the hippocampus and cortical grey matter so do the three weakest; the weakest omits the slowest-clearing member of that cohort, whose removal raises the cohort mean and leaves $P(\mathrm{SD} < \mathrm{R}) = 0.982$, 0.956 and 0.929 in the three regions. By comparison, deletions from the reference cohort that most weaken the contrast leave the probability at 0.988, 0.969 and 0.944.

The model's return-to-baseline assumption was checked against the data. Nineteen of the 21 subjects had a scan roughly four weeks after injection (696–699 h). Tracer had largely cleared in the majority of subjects. The signal had returned close to its pre-injection level with a median enhancement at four weeks of 3% above baseline, half the measurements lying between

2% below and 7% above it, and 88.7% of measurements were within 10% of the baseline value of 1. Residual elevation was concentrated in two subjects, one in each cohort, rather than spread across the group; their median four-week enhancement was 15% and 10% above baseline, elevated in all 42 regions and 41 of 42 regions respectively. This is consistent with genuinely slower whole-brain clearance in these individuals, and with reports of prolonged gadolinium presence following intrathecal administration[5]. Excluding them, the scatter of four-week enhancement about unity has a standard deviation of 5.6%, which is the empirical anchor for the observation-scale prior (Section S5).

These two subjects are not, however, slow clearers in the regions on which the claims rest. The sleep-deprived subject has the fastest fitted clearance of the seven sleep-deprived subjects in all three primary regions, 14 to 72% above that cohort's median, and the reference subject lies within its cohort's range in every region. Retaining tracer at four weeks is thus a whole-brain feature that does not show up as slow clearance region by region. This is because the clearance rate is set by the 24–48 h decline, which a small residual persisting to four weeks does not alter. Removing either subject leaves every conclusion unchanged.

**Table S5. Leave-one-subject-out robustness of the between-cohort contrast.** Each region was refitted with every contributing subject removed in turn (21 refits per region). $\Delta = \mu^{\mathrm{SD}}_{\log k_{\mathrm{out}}} - \mu^{\mathrm{ref}}_{\log k_{\mathrm{out}}}$, so negative values indicate reduced clearance under sleep deprivation. Two deletions are shown individually, being those that most weaken the contrast from each side: omitting the slowest-clearing subject of the sleep-deprived cohort, which raises that cohort's mean, and omitting the fastest-clearing subject of the reference cohort, which lowers the reference mean. The former is the weakest of all 21 refits in every region. SD denotes sleep-deprived and R reference.

| Region | Fit | Median Δ | $P(\mathrm{SD} < \mathrm{R})$ |
|---|---|---|---|
| Entorhinal cortex | full data | −0.414 | 0.994 |
| | omit slowest sleep-deprived | −0.353 | 0.982 |
| | omit fastest reference | −0.373 | 0.988 |
| | range across 21 refits | −0.461 to −0.353 | 0.982–1.000 |
| Hippocampus | full data | −0.389 | 0.980 |
| | omit slowest sleep-deprived | −0.328 | 0.956 |
| | omit fastest reference | −0.342 | 0.969 |
| | range across 21 refits | −0.484 to −0.328 | 0.956–0.995 |
| Cerebral cortex (grey matter) | full data | −0.392 | 0.962 |
| | omit slowest sleep-deprived | −0.336 | 0.929 |
| | omit fastest reference | −0.358 | 0.944 |
| | range across 21 refits | −0.449 to −0.336 | 0.929–0.973 |

### S12 Simulation-based validation: parameter recovery and calibration

We validated the fitting pipeline by simulation in two ways. First, parameter recovery: we generated synthetic data with known clearance rates, using this study's own subjects, acquisition times and noise level, fitted them exactly as the real data, and asked whether the known rates were recovered and their intervals contained them. Second, simulation-based calibration: we drew parameters from the priors themselves, generated data from each draw, refitted, and checked that the true values fell uniformly across their posteriors over the whole range of behaviour the priors allow.

**Design-matched recovery.** Each synthetic cohort was a copy of a real one. For each region, curves were generated at the observed scan times, for the same subjects in the same cohorts, and with the same per-point measurement errors. We used each region's own fitted posterior medians for the reference cohort, noise scale and between-subject spreads as the true parameters, and the study's overall cohort contrast (−0.381), as the true difference in every region. These simulations therefore sit where the real analysis sits. We did this for all 42 regions, 12 times each. We also added two scenarios that moved the truth away from the fitted values: one with no cohort difference at all, in which the sleep-deprived cohort's mean was replaced by the reference cohort's, and another scenario with fitted clearance rates multiplied by 1.5. Those two scenarios were also repeated under a second generator in which amplitude is derived as $A = \kappa C_0/(k_{\mathrm{in}} - k_{\mathrm{out}})$ rather than drawn independently, so that amplitude and rate gap are linked as the mechanism implies (Section S4).

Recovery was scored two ways. First, against the mean of the true parameter values of the subjects drawn in each replicate, which measures error the pipeline itself introduces. Second, against the true cohort mean, from which each replicate's subjects were drawn: this additionally includes the variation from one sample of 14 and 7 subjects to the next, and so gives the precision a study of this size can reach even with a perfect pipeline.

Whether a region recovers the synthetic truth depends on how much signal its curves carry. We measure this as the peak enhancement divided by the observation scale $\sigma_{obs}$ (Fig. S3). Above a ratio of about 2, the 39 regions in that range have a mean absolute bias in cohort clearance of 6.4% and credible-interval coverage of 97%, with no trend from a ratio of 2.5 up to 14. Below 2 lie three deep grey structures, putamen, caudate and pallidum, whose peak enhancement is no larger than their noise. These are biased +11 to +36% toward the prior, and their credible intervals cover the truth only 78% of the time pooled over both cohorts' rates (89% for the reference cohort alone; 50-58% for the sleep-deprived cohort in the putamen and pallidum). Because both cohorts are pulled toward the same prior centre, the difference between them shrinks. These are the same three prior-dependent regions identified by the contraction analysis (Section S7) and by perturbing the priors (Section S9).

In the well-identified range the bias is not centred on zero but sits slightly negative (mean −5.7%), because partial pooling draws the cohort means toward the prior. Cortical clearance rates are therefore likely to be marginally underestimated. The cohort contrast is attenuated in the same direction: the recovered $\Delta$ lies on average 0.06 log units closer to zero than the truth, so a true reduction of 32% is recovered as about 27%, and the reported reduction is if anything conservative; its credible interval contained the true contrast in 95% of replicates. Lateral occipital cortex is the other exception: at a ratio of 2.5 it is the only region above the threshold with a substantial positive bias (+10%), which fits its being the weakest-enhancing cortical region.

The two displaced scenarios, generated from the fusiform gyrus's acquisition times, subjects and fitted noise (a mid-signal cortical region) under both generators, behaved as required. With no cohort difference present, the recovered contrast was −0.08 and +0.02 under the independent and linked generators respectively, with the credible interval covering zero in every replicate; with clearance multiplied by 1.5, the shifted rate was recovered to within 1% and 6%. Deriving the amplitude from the rate gap rather than drawing it independently changed the recovered contrast by 0.09 and 0.11 log units in the two scenarios, within the replicate scatter, and altered no conclusion. No replicate failed to converge in any of the 504 design-matched, or the 48 displaced-scenario fits.

**Simulation-based calibration.** Simulation-based calibration draws every truth from the model's own priors— all ten hyperparameters, with each cohort's mean of the three kinetic parameters drawn separately from the prior the cohorts share — simulates data on the real design, and tests whether the rank of each truth within its own posterior is uniform across replicates. Uniform ranks certify the entire inference machinery, model, weights, sampler and code together, over the full support of the priors and without reference to the fitted values. Over 400 replicates on the precuneus design, 394 converged; the six exceptions were marginal ($\hat{R}$ up to 1.03) and each had at least one cohort drawn with slow clearance, where a single curve carries least information. Rank uniformity held for the reference cohort mean, the sleep-deprived cohort mean and the contrast (Figure S4), with Kolmogorov–Smirnov deviations of 0.052, 0.031 and 0.064 against a 5% critical value of 0.069. The ranks are uniform overall but not independent of the truth: a replicate's rank correlates with its own true value at +0.34, +0.37 and +0.34 for the reference mean, the sleep-deprived mean and the contrast. This follows from the priors being weakly informative rather than flat — a truth from the tail of the prior sits predictably off-centre in its own posterior — and does not affect the uniformity that calibration requires.

**Table S6. Parameter recovery across all 42 regions** (Section S12). Every region was simulated from its own fitted values and design, 12 times each, and refitted. Regions are grouped by the signal their curves carry, taken as peak enhancement divided by the observation scale $\sigma_{obs}$. Bias and RMSE for clearance are percentages of the true rate; those for the cohort difference $\Delta$ are on the log scale, scored against the cohort means actually drawn in each replicate. The final $\Delta$ column is scored instead against the cohort means the simulation was given, and so also carries the variation between one sample of subjects and another. Coverage is of the 95% credible interval for the reference cohort's clearance rate; the sleep-deprived cohort's is given in the text.

| | | clearance $k_{out}$ | | | cohort difference $\Delta$ | | | |
|---|---|---|---|---|---|---|---|---|
| Signal | $n$ | bias | \|bias\| | RMSE | bias | RMSE | vs. given | Coverage |
| below 2 | 3 | +24.6% | 24.6% | 33.5% | +0.29 | 0.43 | 0.44 | 89% |
| 2 to 4 | 4 | −0.3% | 5.5% | 11.0% | +0.07 | 0.18 | 0.25 | 98% |
| 4 to 6 | 8 | −8.8% | 8.8% | 12.2% | +0.06 | 0.15 | 0.19 | 97% |
| 6 or more | 27 | −5.6% | 5.8% | 9.6% | +0.04 | 0.13 | 0.19 | 97% |
| *The three regions below a ratio of* 2, *individually:* | | | | | | | | |
| Putamen (ratio 1.8) | | +36% | | | | | | 83% |
| Pallidum (ratio 0.8) | | +28% | | | | | | 92% |
| Caudate nucleus (ratio 1.3) | | +11% | | | | | | 92% |

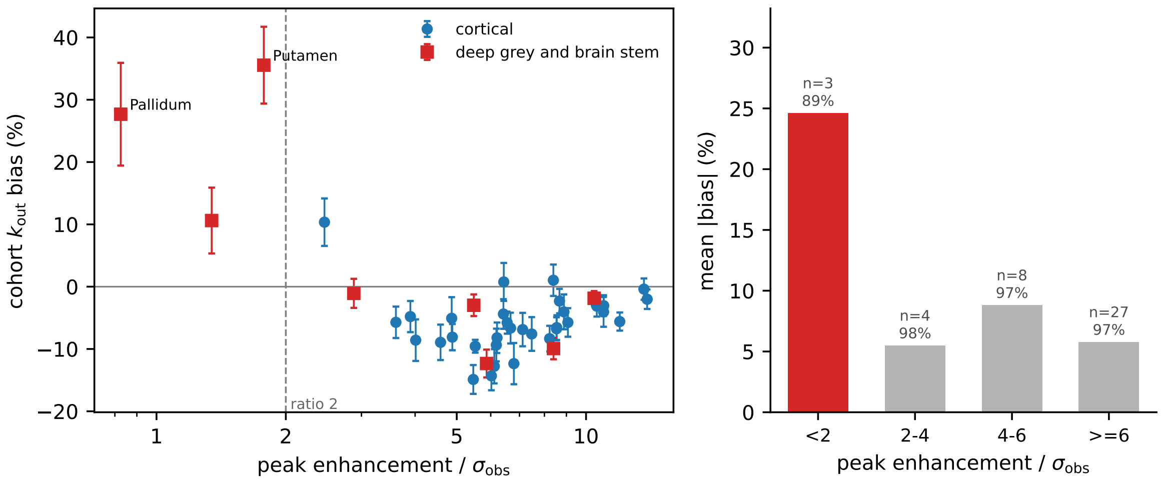


**Figure S3. Recovery depends on the signal a region's curves carry.** Left: bias in recovered reference-cohort clearance for each of the 42 regions, plotted against peak enhancement divided by the observation scale $\sigma_{obs}$, with bars showing the standard error over 12 replicates and the dashed line marking a ratio of 2. Right: mean absolute bias by band, with the number of regions and their mean credible-interval coverage above each bar. Above a ratio of 2 recovery is uniform, with no trend across the range. Below it, the three deep grey structures are biased toward the prior and their credible intervals under-cover.

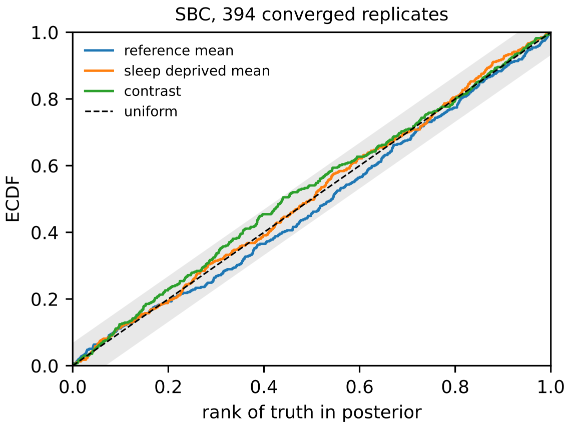


**Figure S4. Simulation-based calibration.** Empirical distribution of the rank of each true value within its own posterior, over 394 converged replicates with all hyperparameters drawn from the priors, for the reference cohort mean, the sleep-deprived cohort mean and their contrast. Correct calibration gives uniform ranks (diagonal); the largest deviation is 0.064, below the 5% critical value of 0.069.

### S13 Consistency of the spatial pattern between subjects

Most of the variation in clearance lies between regions rather than between subjects, and within a cohort subjects differ mainly in their clearance level rather than in the spatial pattern of clearance. Decomposing subject-level log clearance (posterior medians) over the subject-by-region matrix, in the reference cohort 62% of the variance lies between regions, 29% between subjects in overall level, and 9% in subjects departing from the shared regional pattern (Figure S6). In the sleep-deprived cohort the corresponding numbers are 63, 27 and 10%.

The regional ordering of clearance is largely shared between subjects within the same cohort (Figure S5, left) and between cohorts (Figure S5, right). Within the reference cohort a subject's clearance map correlates with the median clearance map of the others at $\rho_s = 0.93$ (range 0.71 to 0.97), and the median correlation between pairs of subjects is 0.87. Within the sleep-deprived cohort a subject's clearance map correlates with the median clearance map of the others at $\rho_s = 0.88$ (range 0.82 to 0.94), and the median correlation between pairs of subjects is 0.80. The two cohort median clearance maps (Figure 2B, C) correlate at

$\rho_s = 0.85$, with medial orbitofrontal cortex fastest in both and entorhinal cortex in the top three of each.

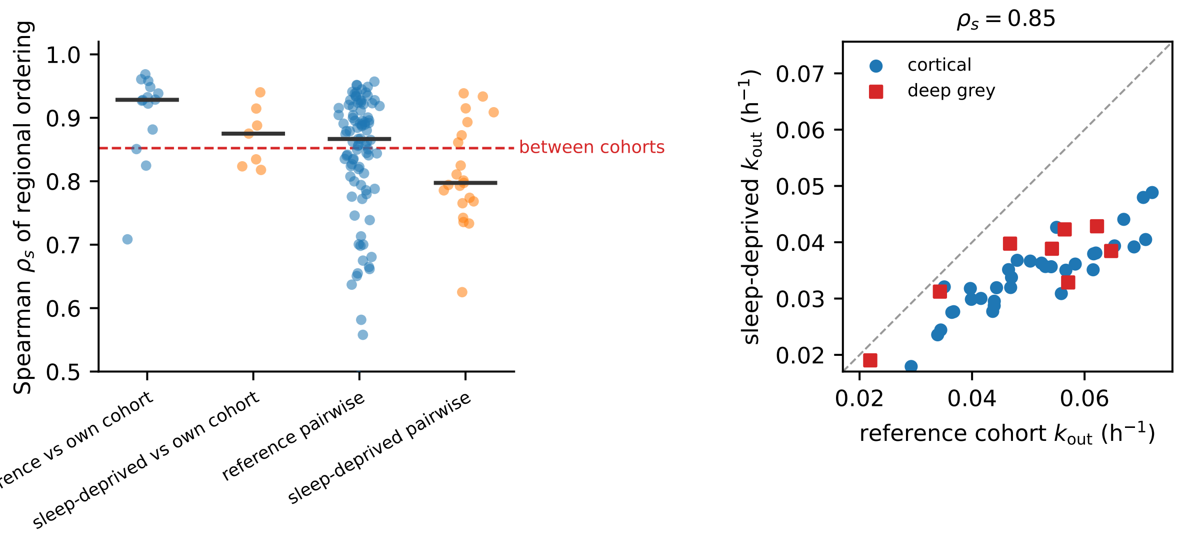


**Figure S5. The regional ordering of clearance is largely shared between subjects.** Left: Spearman correlations between regional orderings across the 42 regions, with bars at the medians. In the first two groups each point is one subject, correlated against the median ordering of the other subjects in the same cohort; in the last two each point is one pair of subjects (91 pairs in the reference cohort, 21 in the sleep-deprived). The dashed line marks the correlation between the two cohorts' median orderings. Right: the clearance rate of each of the 42 regions in the two cohorts, plotted against each other, with the identity line dashed; the title gives the Spearman correlation between them, the same value marked by the dashed line in the left panel.

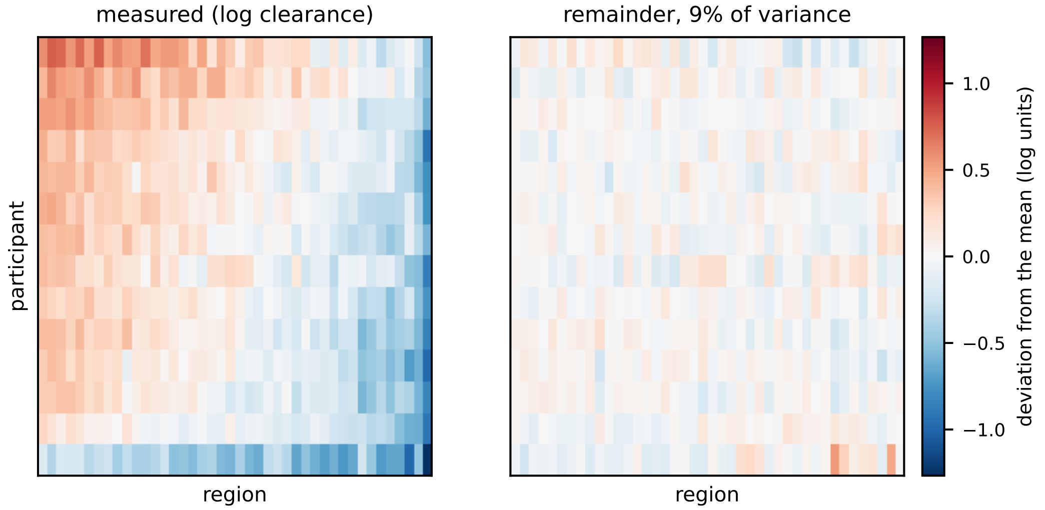


**Figure S6. Subjects differ in how fast they clear, not in which regions clear fastest.** Left: log clearance for the 14 reference subjects across the 42 regions, with the grand mean removed; regions are ordered by their mean rate and subjects by theirs. Right: what remains once each region's typical rate and each subject's overall level are subtracted, on the same colour scale. This remainder is where subject-specific spatial structure would appear, and it accounts for 9% of the variance, against 62% between regions and 29% between subjects.

### S14 Hemispheric structure

Each subject-region was fitted three times: to the left hemisphere measurements alone, to the right alone, and to the mean of the two. The brain stem is midline, leaving 41 paired structures. All comparisons between clearance maps come from the bilateral fits, which draw on both hemispheres. For simulation studies, the hemispheric fits are used to set node-wise clearance in the network model of tau spread, which requires a rate for each hemisphere separately.

Pooling the hemispheres loses no spatial structure. Each hemisphere gives the same ranking of regions by clearance (Spearman $\rho_s = 0.976$, Pearson on the log rates 0.972; Fig. S7), so the pattern reported in the main text does not arise from combining them. The bilateral fit sits between the two hemispheric estimates in 35 of 41 structures, departing from their geometric mean by 0.43% at the median. In the six exceptions the hemispheres almost agree, differing by 0.8% at the median against 3.5% elsewhere, so the gap between them is narrower than the bilateral fit's own variation. Those estimates overshoot the nearer hemisphere by 0.74% at the median and at most 1.5%.

A small asymmetry is present. Structures differ between hemispheres by 3.2% at the median, favouring the right in 26 of 41, with the largest differences in caudal middle frontal cortex (21%), pericalcarine cortex (14%) and the pallidum (12%). It is larger where the fit is less well identified ($\rho_s = -0.30$; Fig. S7). Bilateral fitting averages over it.

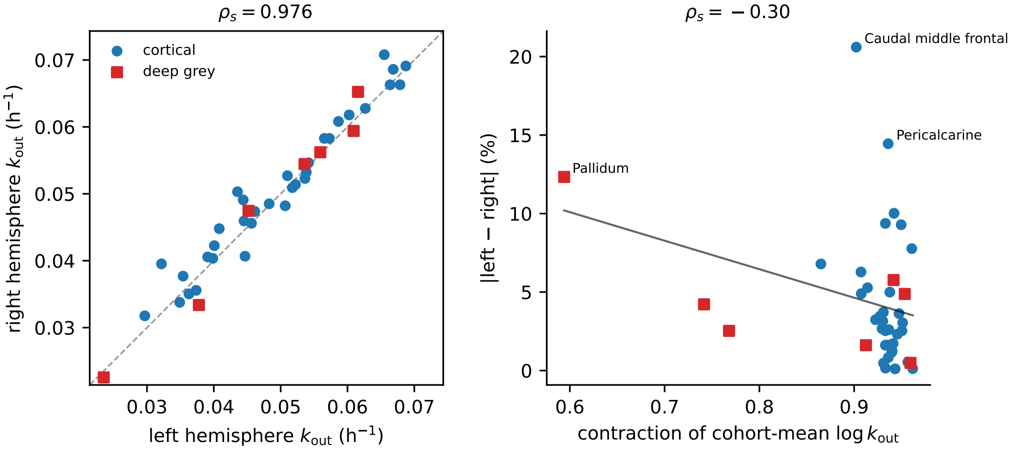


**Figure S7. The two hemispheres give the same regional pattern.** Left: clearance in the right hemisphere against the left, one point per paired structure, with the identity line dashed. Right: the difference between hemispheres against how well each structure's clearance is determined by the data (Section S7), with the three largest differences labelled. Asymmetry is larger where the fit is less well constrained.

### S15 Region size, CSF proximity and partial-volume effects

Regional signal was taken as the median voxel intensity within each parcel. This reduces partial-volume contamination but small parcels bordering CSF spaces remain susceptible. The entorhinal cortex is both small and CSF-adjacent, and we report it among the fastest-clearing regions, so we asked whether the regional pattern is confounded with geometry. Region size was taken as the mean voxel count across subjects, summing hemispheres, and each region's cohort-level clearance rate in the reference cohort was regressed on $\log_{10}$ size across the 42 regions. A region's residual from that line is its clearance relative to what its size predicts. Rank correlations are Spearman's $\rho_s$ throughout.

Smaller parcels do clear faster ($\rho_s = -0.46$, $p = 0.002$). Regions bordering a CSF space also clear faster than those that do not: median 0.058 against 0.044 $\mathrm{h}^{-1}$ ($p = 0.0003$). The 16 CSF-adjacent regions were labelled by anatomy: medial temporal, cingulate, medial occipital and insular cortex, the ventricle-lining nuclei (caudate, thalamus, accumbens) and the brain stem. The label was not derived from a measured distance.

Between-cohort contrasts use the same regions in different subjects, so a size effect enters both cohorts and cancels. The size of a region predicts $k_{\mathrm{in}}$ almost as strongly as $k_{\mathrm{out}}$ ($\rho_s = -0.41$ against $-0.46$), so it acts on the whole curve. Under sleep deprivation clearance falls by 31% (95% credible interval 8 to 48%) while influx does not change (+1.5%, −23 to +36%; main text). Size is therefore a curve-wide effect, and the effect of sleep deprivation that we report is not.

Within-cohort comparisons survive adjustment. Entorhinal cortex has the third-largest positive residual from the size regression of the 42 regions ($+0.0129\,\mathrm{h}^{-1}$, 1.3 standard deviations), so it clears faster than its size predicts, and it exceeds each subject's own whole-brain mean in all 21 subjects (median elevation +41%, minimum +20%).

Partial-volume effects bear on only one of the three kinds of comparison we make; between-region comparisons. Size and Braak stage are largely unrelated in this parcellation ($\rho_s = +0.29$, $p = 0.08$), so the decline of clearance across Braak stages is the same whether computed on the raw rates or on the size residuals ($\rho_s = -0.42$ raw against $-0.41$ adjusted, $n = 36$). The limbic lobe remains the fastest-clearing lobe on the same basis, its regions clearing $+0.0099\,\mathrm{h}^{-1}$ faster at the median than their size predicts, against $-0.0005$ to $-0.0064$ for the other four.

We report unadjusted rates throughout the main text, since this is a sensitivity check rather than a correction and residuals have no physical meaning. Partial-volume effects are therefore present in these data but do not account for what we report. Parcel size predicts clearance, and it does so through the whole enhancement curve rather than through efflux alone. Neither the entorhinal elevation nor the Braak gradient survives only because of it: both results hold once size is accounted for.

One consequence of smaller parcels clearing faster is that regional summaries depend on how regions are weighted. Means across regions are taken unweighted when analysing the clearance maps; weighting instead by parcel volume lowers the whole-brain mean by 8.3% and raises the entorhinal margin from +41 to +53%, because the largest parcels (superior frontal, rostral middle frontal, superior and inferior parietal cortex) are slow-clearing. The unweighted convention is the conservative one and treats each region as a single observation, which is what the regional comparisons ask about.

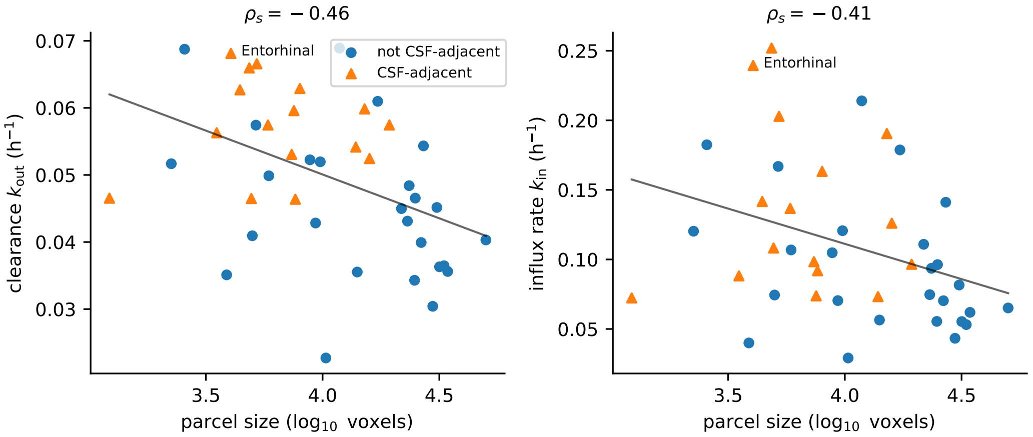


**Figure S8. Parcel size predicts both clearance and influx.** Cohort-level clearance rate (left) and enhancement rate (right) of the reference cohort against parcel size, one point per region, with the least-squares line; CSF-adjacent regions as triangles, the entorhinal cortex labelled. Size is negatively related to both rates ($\rho_s = -0.46$ and $-0.41$), so it acts on the whole enhancement curve, whereas sleep deprivation lowers clearance without changing influx. The entorhinal cortex lies well above the line: it clears faster than its size predicts.

### S16 Network model graph, parameters and their sensitivity

Simulations were performed on the same high-resolution structural connectome used by Putra *et al.*[6], corresponding to Scale-500 of the Lausanne multiresolution atlas[7]. The network has 1015 nodes, and was constructed from Human Connectome Project data using MRtrix deterministic tractography with 20,000 streamlines and randomised seeding[8]. Edges are weighted by $n_{ij}/l_{ij}$, the mean number of fibres between two nodes divided by their mean length (Methods). These weights are divided by a single constant, chosen so that the mean row sum of the adjacency matrix matches the $n_{ij}/l_{ij}^2$ weighting used in the earlier model[9]. The Laplacian is therefore dimensionless, and $\rho$ is a rate in inverse years.

For each subject $s$, we estimated hemispheric clearance rate $k^s_{\text{out},r}$ for each region $r$ of the 83-region parcellation. To construct the initial clearance field on the $N = 1015$-node simulation network, each high-resolution node $i$ is assigned the measured clearance rate of its parent region, denoted $r(i)$. The imaging-derived clearance map therefore enters the propagation model as the initial condition

$$\lambda_i^s(0) = \gamma k^s_{\text{out},r(i)}, \qquad i = 1,\ldots,N, \tag{8}$$

where the same scaling constant $\gamma$ is applied to every node and every subject. Thus, the spatial pattern and all relative differences in clearance between regions, subjects and cohorts are preserved, while the absolute clearance scale is calibrated to that of the propagation model. The protein field is initialised separately, with non-zero $p_i(0)$ confined to nodes belonging to the entorhinal cortex, as described in the main Methods. The measured clearance maps therefore determine the initial clearance field $\lambda(0)$, whereas the entorhinal seed determines the initial protein field $p(0)$.

The parameters for the propagation model Eq. [1] (main text)were adopted from prior work rather than estimated here: $\rho$, $\alpha$, $\beta$ and $\lambda_{\text{crit}}$ are as in [9]. Parameter values and their derivation are detailed in Table S7. The one quantity calibrated in this work is the common scaling factor $\gamma$ in Equation (8). Because gadobutrol clearance is used here as a proxy for τ clearance, the absolute rates need not coincide. The factor $\gamma$ therefore maps the measured tracer-clearance rates onto the clearance scale of the propagation model while preserving their relative differences across regions, subjects, and cohorts. It is chosen so that the reference cohort's mean initial clearance is $0.40\,\lambda_{\text{crit}}$:

$$\gamma = \frac{0.40\,\lambda_{\text{crit}}}{\bar{k}_{\text{out}}}, \qquad \bar{k}_{\text{out}} = \frac{1}{14}\sum_{s=1}^{14}\frac{1}{N}\sum_{i=1}^{N} k^{s}_{\text{out},r(i)}, \tag{9}$$

where $\bar{k}_{\text{out}}$ is the reference cohort's mean measured clearance, obtained by averaging each subject's clearance map over the $N = 1{,}015$ network nodes and then over the 14 reference subjects. This gives a reference-cohort mean initial model clearance of $\bar{\lambda}(0) = \gamma\bar{k}_{\text{out}} = 0.40\,\lambda_{\text{crit}}$. With $\lambda_{\text{crit}} = 0.72\,\text{yr}^{-1}$ and $\bar{k}_{\text{out}} = 0.047\,\text{h}^{-1}$, $\gamma = 6.13\,\text{h}, \text{yr}^{-1}$. The adopted calibration yields simulated pathology progressing over several decades, consistent with the timescale of Alzheimer's disease; sensitivity to this choice is examined below.

**Table S7. Network model parameters** Propagation-model parameters follow prior work where indicated; the calibration factor $\gamma$ is determined in this work. Sensitivity to each is reported in this section.

| Parameter | Value | Basis for the value |
|---|---|---|
| $\rho$ | $1 \times 10^{-2}$ yr$^{-1}$ | Chosen with $\alpha$ so that the maximal rate of toxic protein increase matches modelling studies fitted to Alzheimer's imaging data[10–12] |
| $\alpha$ | 2.1 yr$^{-1}$ | As above |
| $\beta$ | 1.0 yr$^{-1}$ | Consistent with a typical disease duration of about 30 years[9] |
| $\lambda_{\text{crit}}$ | 0.72 yr$^{-1}$ | Critical clearance scale derived from protein-aggregation kinetics[13] |
| $\lambda_i^{\infty}$ | $1 \times 10^{-6}$ yr$^{-1}$ | A low but non-zero floor on regional clearance, following the model structure of[9] |
| $\gamma$ | 6.13 h yr$^{-1}$ | Calibration of tracer-derived clearance to the model clearance scale (this work) |

We varied each parameter in turn and repeated every simulation, recording the simulated onset times of Braak I and Braak VI, the correlation between a subject's mean clearance and their time to Braak VI, and the shift in Braak I onset when the clearance maps were homogenised (Fig. S9). The qualitative conclusions were robust across the parameter ranges examined. Across 23 alternative settings, the correlation remained between 0.97 and 0.99, compared with 0.98 in the main analysis, and homogenising the maps advanced Braak I in every setting, by 1.1–6.5 yr compared with 3.2 yr in the main analysis. The onset times changed as expected but generally modestly: doubling $\rho$ speeds propagation and shortens the Braak I-to-VI interval by 7%, while varying $\alpha$ and $\beta$ over the same relative range changes it by no more than 4%.

$\lambda_{\text{crit}}$ requires separate treatment, because the clearance maps are rescaled so that the reference cohort's mean sits at a fixed fraction of it. Varying it with that fraction held constant rescales time and nothing else: halving it lengthens the Braak I to VI interval by 92% and doubling it shortens it by 48%, while leaving the correlation and the direction of the flattening effect untouched. It sets the timescale on which the simulation runs rather than the behaviour it exhibits.

The final column of Fig. S9 shows the sensitivity to the calibration constant $\gamma$. Since $\gamma$ is fixed by where it places the reference cohort's mean clearance relative to the critical rate, the horizontal axis is that ratio, $\bar{\lambda}(0)/\lambda_{\text{crit}}$, with the adopted value 0.40. Raising it from 0.25 to 0.50 increases the advance in Braak I onset on flattening the maps from 1.1 to 6.5 yr. This is because the further a cohort's clearance level sits below the critical rate, the more protective its spatial arrangement is to simulated τ seeded in the entorhinal cortex. This is the same dependence that makes the entorhinal protection smaller in the sleep-deprived cohort, appearing here as a property of the model rather than of the measured maps. At the adopted value of 0.40 the fastest-clearing region is $0.90\,\lambda_{\text{crit}}$; by 0.45 it crosses the threshold, beyond which that region would cease to accumulate protein at all, so the calibration places the cohort close to but below a regime change in the model.

### S17 Does the arrangement of clearance matter, or only its spread?

The homogenisation analysis (Results) flattens each map toward its mean, which removes the regional spread and the arrangement of rates together. To separate them, each reference subject's 83 per-hemisphere regional clearance rates were randomly permuted across regions: the distribution is preserved exactly—the same mean, spread and extremes, the same 83 numbers—and only which region holds which rate changes. Permutation acts at the region level, since clearance was measured per region and permuting nodes within a region would create structure the data never resolved. Each subject's measured map was compared against 1,000 permutations of their own values, simulated on their own network with the same model, seed and threshold as the main analysis (Figure S10).

The measured arrangement delays initiation and compresses the course. Relative to the mean of its own permutations, the measured map reached Braak I later in all 14 subjects (median +2.9 yr), lying above 98% of its own permutations at the

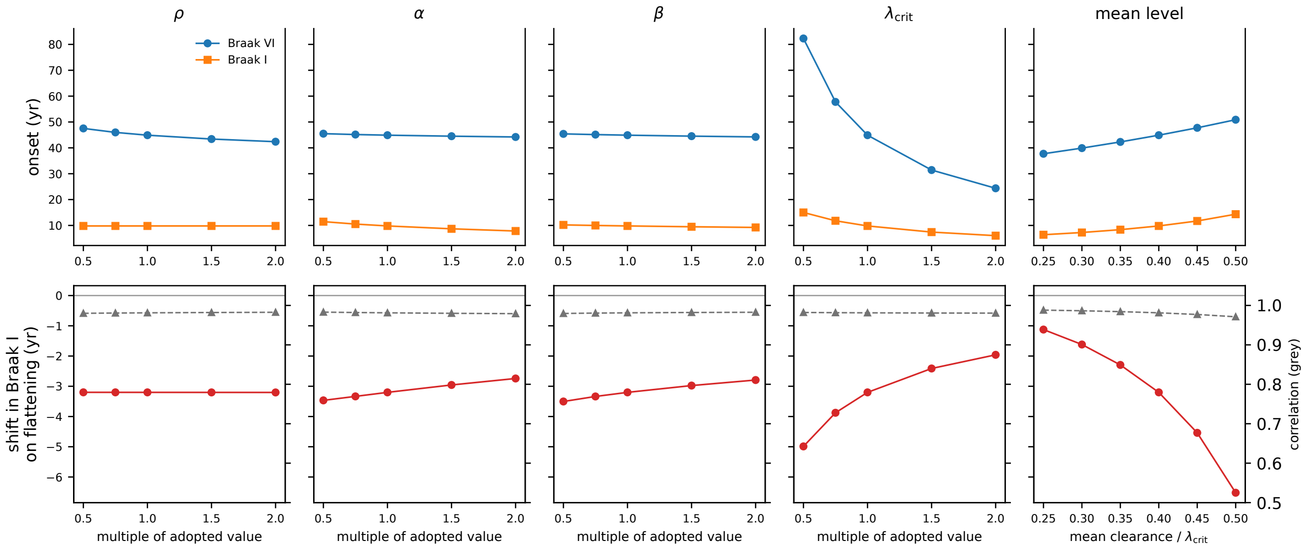


**Figure S9. Sensitivity of the simulation to the model's parameters.** Each column varies one parameter about its adopted value, the first four as a multiple of it and the last as the fraction of $\lambda_{\text{crit}}$ at which the reference cohort's mean clearance is set. Top row: median onset of Braak stages I and VI across the 14 reference subjects. Bottom row: the two results the paper's claims rest on, the shift in Braak I onset when the clearance maps are flattened at matched level (red, left axis) and the correlation between a subject's mean clearance and their time to Braak VI (grey, right axis). The correlation is unchanged throughout, and flattening advances Braak I in every setting.

median subject (range 91–100%), and reached Braak II later in all 14 (+6.9 yr) and Braak III later in 13 of 14 (+3.1 yr). From Braak IV the pattern reverses: the measured map was neutral at stage IV (later in 4 of 14, median −0.4 yr) and reached stages V and VI earlier than its permutations on average (later in 0 and 1 of 14 respectively; −1.7 and −2.0 yr). This is the same compression the homogenisation analysis produced, now against rearrangements of the same rates rather than against a flattened map. The gain at Braak I ranged from under a year to twelve years across subjects, and was larger in subjects whose pathology begins later ($\rho_s = +0.96$ between measured onset and the gain over permutations).

The permutations also locate the effect at the seed. Across all permutations, pooled over subjects, the clearance a permutation happened to place at the entorhinal cortex predicts the Braak I onset it produces ($r = +0.93$; Figure S10, right), whereas it predicts the later stages only weakly ($r = 0.09$–$0.24$). Permutations that placed a rate in the top quarter of the subject's own values at the seed delayed Braak I by 2.0 yr relative to the rest, and those placing a rate at least as high as the measured entorhinal rate by 3.5 yr. The measured maps sit at the fast end of that relation in every subject.

**Permuting bilateral structures as units.** Permuting the 83 hemispheric regions independently breaks the measured left–right symmetry of the maps (Section S14), which the real arrangement possesses and almost no permutation does. As a stricter null, we therefore repeated the test permuting the 42 bilateral structures as units, each left–right pair moving together with its measured values, so that every permutation retains the same symmetry as the measured map (Figure S11). The result is the same in direction and size and, as expected of a null that shares more structure with the data, less extreme in rank: the measured map reached Braak I later than its permutations in all 14 subjects (median +2.7 yr), lying above 92% of them at the median subject (range 84–98%); Braak VI arrived 2.3 yr earlier and the Braak I–VI interval was 4.8 yr shorter. Permutations placing a top-quarter rate at the seed delayed Braak I by 2.8 yr relative to the rest, and those placing at least the measured entorhinal rate by 3.2 yr.

Together, the two nulls show that the arrangement matters independently of the spread, that it acts through what it places at the seed, and that the arrangement estimated from reference subjects delays initiation more than the great majority of other rearrangements of the same clearance rates.

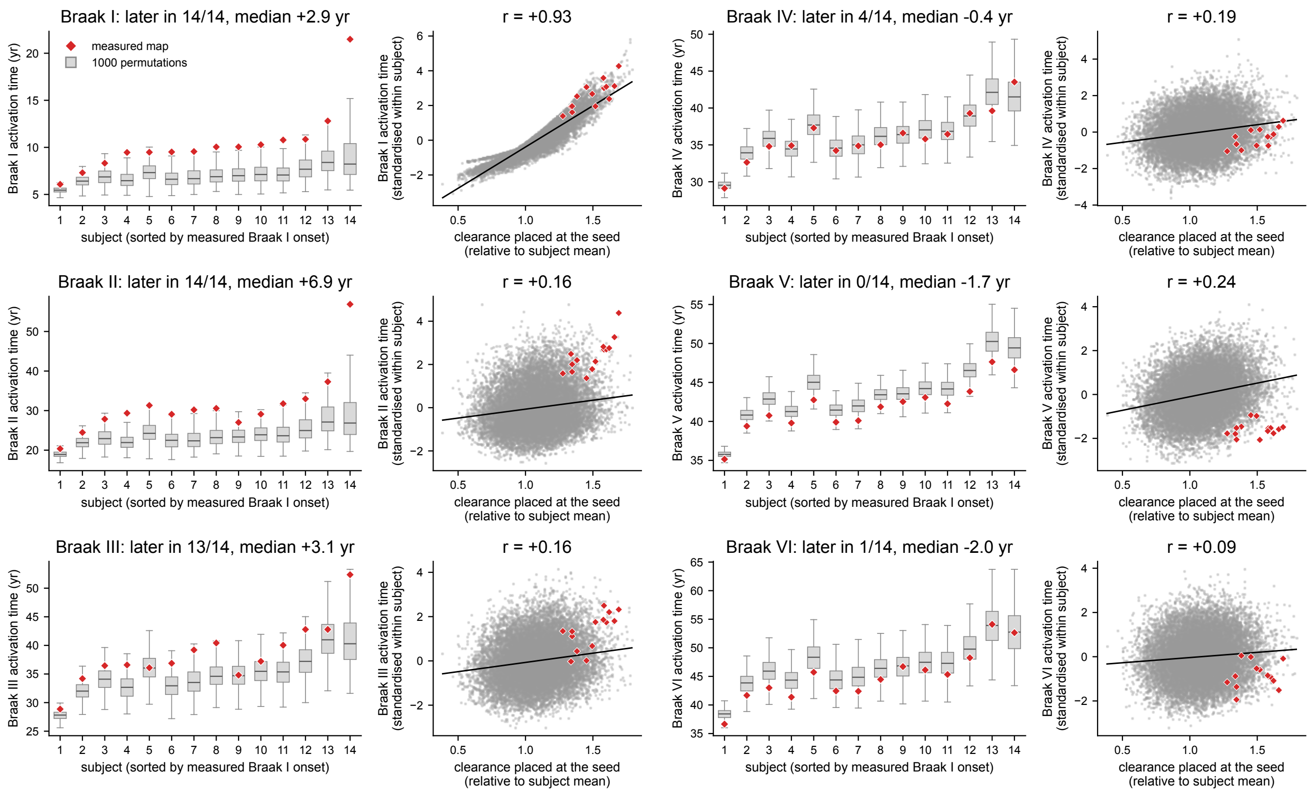


**Figure S10. Permuting per-hemisphere clearance rates across regions.** Each reference subject's 83 per hemisphere regional clearance rates were permuted across regions 1,000 times, preserving their distribution exactly, and each permutation was simulated. Left block, Braak I–III; right block, Braak IV–VI; within each stage, the left panel shows activation time under the permutations (box; whiskers to the range) and under the measured map (diamond), subjects sorted by measured Braak I onset in every panel, and the right panel shows, within the permutations, the clearance a permutation placed at the entorhinal seed relative to the subject's mean against the activation time it produced, standardised within subject and pooled over the 14 subjects, with the measured maps as diamonds and a least-squares line.

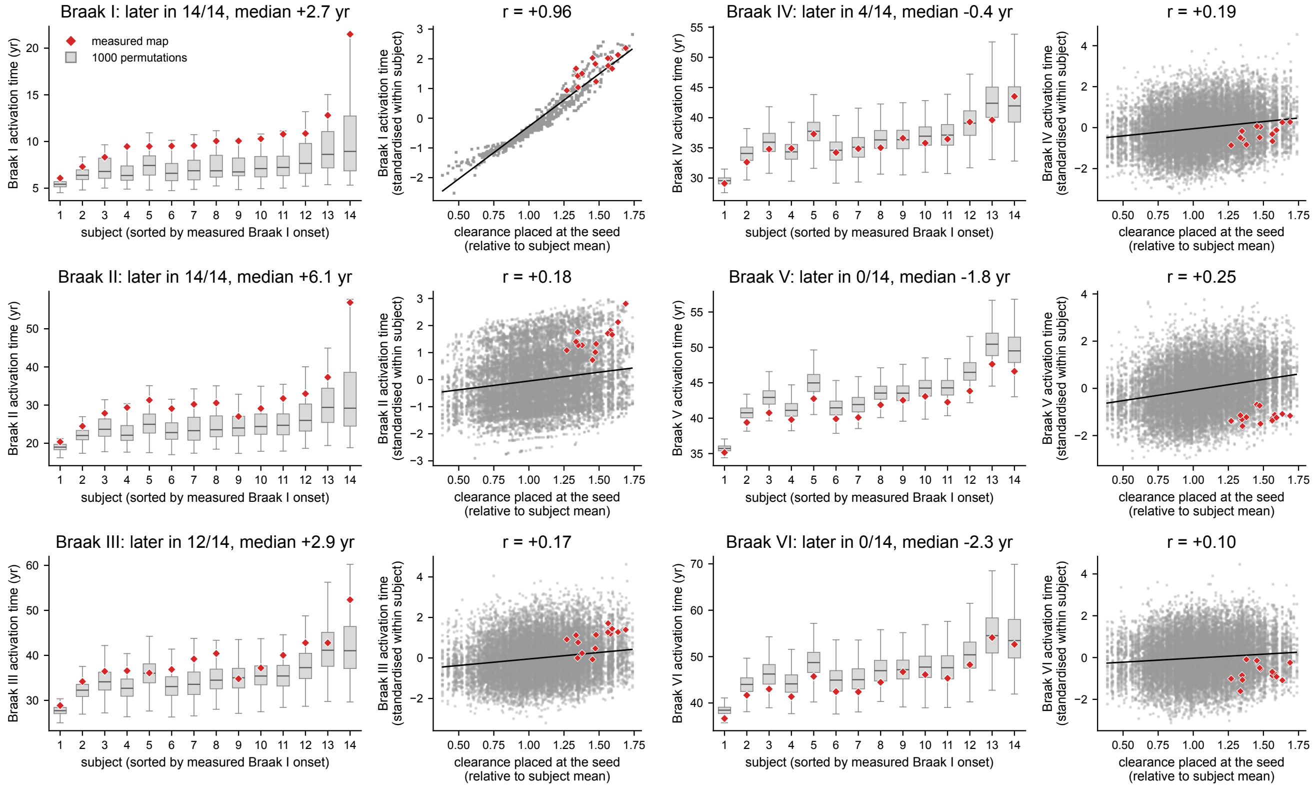


**Figure S11. Permuting bilateral clearance rates across regions.** Each reference subject's 42 bilateral regional clearance rates were permuted across regions 1,000 times, preserving their distribution exactly, and each permutation was simulated. Left block, Braak I–III; right block, Braak IV–VI; within each stage, the left panel shows activation time under the permutations (box; whiskers to the range) and under the measured map (diamond), subjects sorted by measured Braak I onset in every panel, and the right panel shows, within the permutations, the clearance a permutation placed at the entorhinal seed relative to the subject's mean against the activation time it produced, standardised within subject and pooled over the 14 subjects, with the measured maps as diamonds and a least-squares line.
